# *ESR1* fusion proteins invoke breast cancer subtype-dependent enrichment of ligand independent pro-oncogenic signatures and phenotypes

**DOI:** 10.1101/2023.09.18.558175

**Authors:** Megan E. Yates, Zheqi Li, Yiting Li, Hannah Guzolik, Xiaosong Wang, Tiantong Liu, Jagmohan Hooda, Jennifer M. Atkinson, Adrian V. Lee, Steffi Oesterreich

## Abstract

Breast cancer is a leading cause of female mortality and despite advancements in diagnostics and personalized therapeutics, metastatic disease largely remains incurable due to drug resistance. Fortunately, identification of mechanisms of therapeutic resistance have rapidly transformed our understanding of cancer evasion and is enabling targeted treatment regimens. When the druggable estrogen receptor (ER, *ESR1*), expressed in two-thirds of all breast cancer, is exposed to endocrine therapy, there is risk of somatic mutation development in approximately 30% of cases and subsequent treatment resistance. A more recently discovered mechanism of ER mediated endocrine resistance is the expression of ER fusion proteins. ER fusions, which retain the protein’s DNA binding domain, harbor *ESR1* exons 1-6 fused to an in-frame gene partner resulting in loss of the 3’ ER ligand binding domain (LBD). In this report we demonstrate that in no-special type (NST) and invasive lobular carcinoma (ILC) cell line models, ER fusion proteins exhibit robust hyperactivation of canonical ER signaling pathways independent of the ligand estradiol or anti-endocrine therapies such as Fulvestrant and Tamoxifen. We employ cell line models stably overexpressing ER fusion proteins with concurrent endogenous ER knockdown to minimize the influence of endogenous wildtype ER. Cell lines exhibited shared transcriptomic enrichment in pathways known to be drivers of metastatic disease, notably the MYC pathway. The heterogeneous 3’ fusion partners, particularly transcription factors *SOX9* and *YAP1*, evoked varying degrees of transcriptomic and cistromic activity that translated into unique phenotypic readouts. Herein we report that cell line activity is subtype-, fusion-, and assay-specific suggesting that the loss of the LBD, the 3’ fusion partner, and the cellular landscape all influence fusion activity. Therefore, it will be critical to generate additional data on frequency of the ER fusions, in the context of the clinicopathological features of the tumor.

**Significance:** ER fusion proteins exhibit diverse mechanisms of endocrine resistance in breast cancer cell lines representing the no special type (NST) and invasive lobular cancer (ILC) subtypes. Our emphasize upon both the shared and unique cellular adaptations imparted by ER fusions offers the foundation for further translational research and clinical decision making.

## Introduction

Breast cancer comprises one-third of all malignancies occurring in women and is estimated to result in a total of 43,700 U.S. deaths in 2023 alone, statistics which reinforce breast cancer’s status as a modern epidemic (1, 2). Recent advancement in tumor molecular subtyping has provided insights into oncological management and therapeutic intervention (3). A predominant breast cancer molecular classification, the luminal subtype, can be diagnosed by expression of estrogen receptor alpha (ER*α*, *ESR1*; herein denoted ER), a protein which is expressed in 70% of all breast cancers. ER is a hormone receptor and estrogen (E2)-dependent transcription factor of survival and proliferation pathways critical for mammary gland development. Pathway dysregulation as a result of ER overexpression in epithelial cells promotes carcinogenesis (4, 5). Hormonal therapies, including aromatase inhibitors (i.e. anastrozole and letrozole) and selective estrogen receptor modulators (SERMs), notably tamoxifen, yield a 40% reduction in mortality of primary breast cancer (6). The success of hormonal therapy, however, is limited; approximately 30% of early breast cancer cases develop *de novo* or acquired resistance, an additional 70% of patients with metastatic disease lack an appreciable treatment response (7–12). Identifying therapeutic response and investigating mechanisms of endocrine resistance remain a prominent research endeavor in this global health crisis crusade.

Acquired endocrine resistance to selective hormonal therapies occurs through a multitude of adaptations. The most notable resistance mechanism involves genetic alterations in *ESR1*. Mutations in the ligand binding domain (LBD) of ER induce a ligand-independent and constitutively active ER protein which is found enriched in metastatic, endocrine resistant disease resulting in poor prognosis (13, 14). ER amplification has also been suggested as a potential mechanism of tumor evasiveness and survival (7, 15). Interestingly, ER loss is not believed to be a predominant mode of endocrine resistance as the majority of resistant tumors retain their ER positivity (16). In the last decade, enrichment of ER fusion proteins in metastatic tumors that have been treated with endocrine therapy have been identified as a novel mechanism of endocrine resistance (17, 18).

Malignancy associated fusion proteins have been studied for over half a century since the discovery of the recurrent Philadelphia chromosome in chronic myeloid leukemia (19). Identification of disease specific balanced chromosomal rearrangements, and in particular translocations, has resulted in clinical and pharmaceutical advancements, particularity in hematological malignancies, as well as a subset of solid tumors, including prostate and non-small-cell lung cancer (19, 20). Work from The Cancer Genome Atlas (TCGA) revealed an abundance of gene fusions in breast, ovarian, and stomach cancers (21). Gene fusion products resulting in constitutive oncogene activation or tumor suppressor gene silencing have been described as primary drivers of numerous cancer types (20). A crucial caveat to gene fusion discovery, however, is the actionability of identified in-frame fusion transcripts and their subsequent potential translational implications (22).

Gene fusion events involving *ESR1* in breast cancer have been previously described, with one of the most common being a recurrent rearrangement between *ESR1* (Chromosome 6q) exons 1-2 and its neighboring gene, coiled-coil domain containing 170 (ESR1-CCDC170) (23, 24). Identification of genomic structural rearrangements in *ESR1* in metastatic breast cancer unveiled recurrent fusion transcripts at an *ESR1* intron 6 translocation breakpoint (17). Fused to heterogeneous 3’ partner genes, these *ESR1* fusion transcripts retain exons 1-6, but miss their 3’ ER LBD and thus were hypothesized to be ligand independent. Targeted sequencing uncovered interchromosomal *ESR1* fusions to the disabled homolog 2 gene (Chromosome 5p; ESR1-DAB2) identified in a lymph node metastasis, and to the SRY-box transcription factor 9 gene (Chromosome 17q; ESR1-SOX9) discovered in a solid metastatic liver lesion (17). Both of these fusions were discovered in patient tumors that had been previously treated with hormonal therapies. An ER fusion to the lipoma preferred partner (Chromosome 3q; ESR1-LPP) was identified in a patient derived xenograft (PDX) model in collaboration with Champions Oncology (Champions TumorGraft (CTG) CTG-1350); the xenograft was derived from a metastatic liver lesion. In addition, an *ESR1* fusion to the Yes1 associated transcriptional regulator functional protein domains (Chromosome 11q, ESR1-YAP1) was also described in a PDX model (25). ESR1-DAB2, ESR1-SOX9 and ESR1-YAP1 overexpression in a cell line model revealed ligand independent and endocrine resistant fusion activity (17, 18). In the T-47D no specific type (NST; historically referred to as invasive ductal carcinoma, IDC) cell line, overexpression of a subset of identified *ESR1* fusion proteins revealed an estrogen response signature that was subsequently developed into a 24 gene signature as a proxy for functional fusion activity (26). To date, this has been the only study to functionally characterize a set of identified *ESR1* fusion proteins, albeit using only one histological subtype of breast cancer. To further evaluate these ER chimeric proteins, we have engineered NST and invasive lobular carcinoma (ILC) models, depleted of endogenous wildtype ER, to assess fusion functionality in unique cellular backgrounds. Furthermore, we performed multi-omic analyses of NST fusion positive cells to further understand transcriptomic differences among fusions. Evaluation of *ESR1* fusion functionality in various cellular contexts exploits subtype-dependent shared and unique characteristics that a 3’ fusion partner may impart upon one of breast cancer’s most prognostic and clinically actionable genes, *ESR1*.

## Results

### *ESR1* fusions discovered in metastatic disease are hyperactive and ligand-independent when over-expressed in *in vitro* models

Limited previous studies have discovered (**Fig. 1A**) and subsequently analyzed ER fusion proteins (**Supplementary Fig. S1**). Transient expression of the *ESR1* fusion plasmids in human embryonic kidney HEK293 cells demonstrated expression of protein products at anticipated ER wildtype (ESR1-WT), ER truncation (*ESR1* exons 1-6 without ER C’ terminal domain (ΔCTD)), or fusion protein size, detected by both anti-ER*α* and anti-HA antibodies (**Fig. 1B**). Examination of transcriptional activity using an estrogen receptor response element (ERE)-tk-Luc reporter showed that *ESR1* fusions exhibited greater activity in vehicle conditions compared to cells expressing ESR1-WT or ESR1-ΔCTD. Ligand stimulation with 1nM estradiol (E2) resulted in enhanced transcriptional activity of ESR1-WT compared to vehicle conditions (**Fig. 1C**), a phenomenon that was not seen with *ESR1* fusion proteins and ESR1-ΔCTD. Co-treatment with E2 (1nM) and 100nM of either the selective estrogen receptor degrader (SERD) fulvestrant (ICI182-780, ICI) or SERM tamoxifen (Tam) diminished ESR1-WT transcriptional activity compared to vehicle conditions; however, these treatments had no effect on *ESR1* fusion or ESR1-ΔCTD transcriptional activity (**Fig. 1C**). Thus, *ESR1* fusions show ligand-independent hyperactivation.

**Figure 1:**
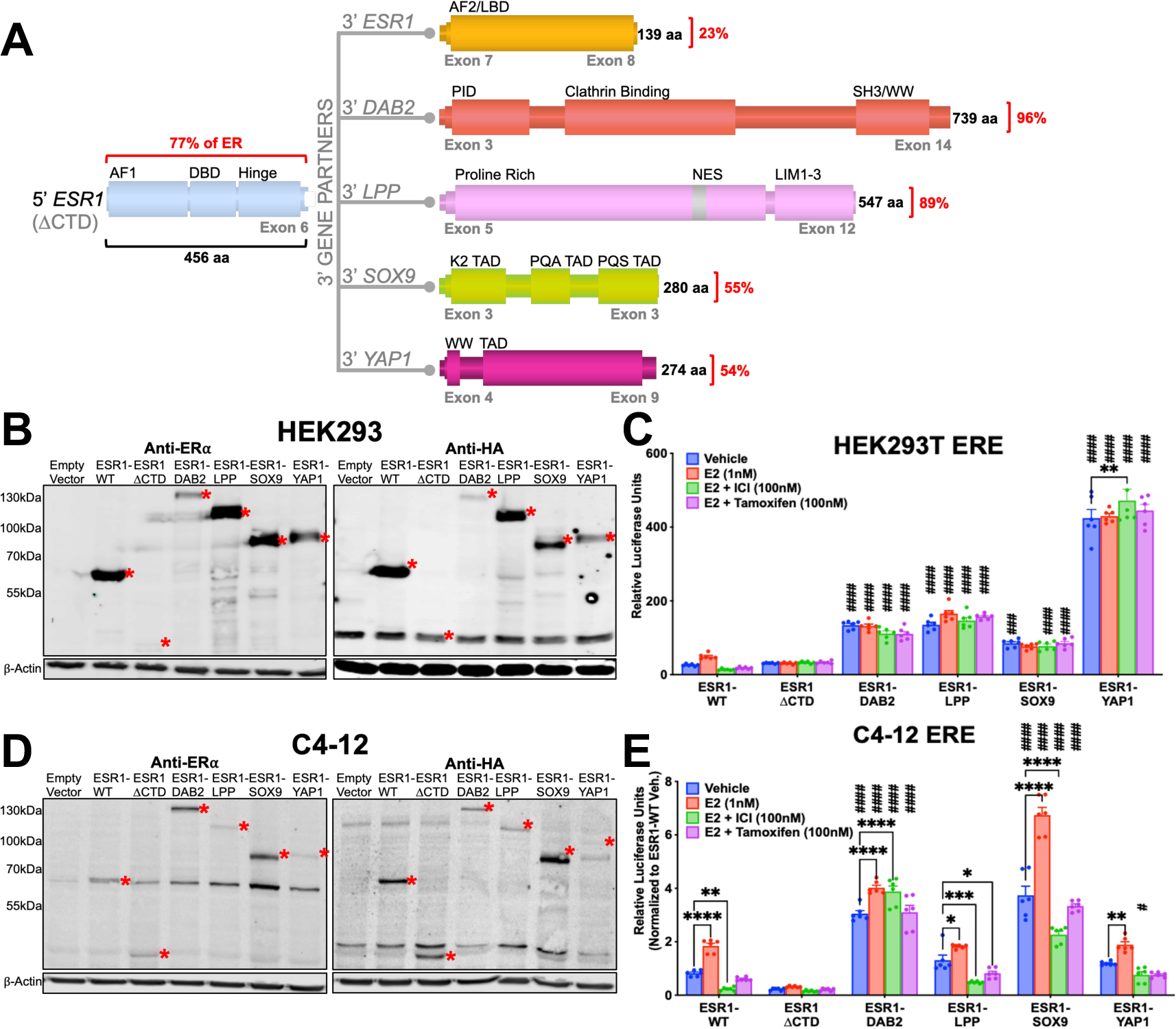
ER Fusions are hyperactive in a ligand resistant manner. (A) Schematic of ER fusions. Red percentages denote quantity of protein fused to the 5’ portion of ER (77% of the ER protein contained within exons 1-6 without C’ terminal domain (ΔCTD)). Activation function (AF), DNA binding domain (DBD), ligand binding domain (LBD), phosphotyrosine interaction domain (PID), SRC homology3 binding domain (SH3), Rsp5 or WWP domain (WW), nuclear export sequence (NES), LIM zinc-binding domain, transcriptional activation domain (TAD), proline-, glutamine-, and alanine-rich (PQA), proline-, glutamine-, and serine-rich (PQS), substrate binding region (Sub), glycogen synthase 1 interaction region (GYS1). (B) Anti-ER and anti-HA immunoblots of transiently overexpressed HA tagged wildtype ER (ESR1-WT), HA tagged truncated ER (ESR1-ΔCTD) and HA tagged ER fusion proteins in HEK293T cells. (C) ERE assay in HEK293T cells transiently expressing ER constructs. Raw luminescence values normalized to internal control renilla luminescence. Representative experiment shown with bar graph equivalent to readings ± SEM, n=2. (D) Anti-ER and anti-HA immunoblots of stably overexpressed ESR1-WT, ESR1-ΔCTD and ER fusion proteins in C4-12 cell line. Construct proteins denoted by red asterisks in all blots. β-actin serves as loading control in all blots. (E) ERE assay in C4-12 cells stably expressing ER constructs. Raw luminescence values normalized to internal control renilla luminescence and ESR1-WT vehicle activity. Representative experiment shown with bar graph equivalent to readings ± SEM, n=3 for each experiment. 2way ANOVA with posthoc Dunnett’s multiple comparisons test showing enrichment in comparison to corresponding ESR1-WT treatment (#) or to intra-construct vehicle treatment (*) in each assay. */#p=0.0332, **/##p=0.0021, ***/###p=0.0002, ****/####p<0.0001.

We next stably expressed the fusion proteins in a breast cancer cell line (C4-12) which was derived from MCF7 cells but has low to absent endogenous ER (**Fig. 1D**) (27). Interestingly, re-expression of endogenous ER was reproducibly observed in cells stably co-expressing *ESR1* fusion proteins (**Fig. 1D**). In an ERE assay, the stable expression of the ESR1-WT plasmid led to an estradiol response which was blocked by fulvestrant and tamoxifen (**Fig. 1E**). ESR1-ΔCTD exhibited minimal activity and there was no effect of E2 or ICI/Tam. ESR1-DAB2 and ESR1-SOX9 demonstrated hyperactivity in the absence of E2 and a minimal response to E2 and ICI/Tam (**Fig. 1E**). ESR1-LPP and ESR1-YAP1 did not demonstrate ligand independent or endocrine resistant activity, mimicking ESR1-WT (**Fig. 1E**).

We sought to evaluate the fusions in NST and ILC cell line models, T-47D and MDA-MB-134-IV (MM134), respectively. To minimize a contribution from endogenously expressed ER, we knocked down ER using a small interfering RNA (siRNA) targeting the 3’ untranslated region of *ESR1*. Transient expression of siESR1 reduced endogenous ER in the T-47D cell line which was most pronounced with transient co-expression of ESR1-SOX9 and ESR1-YAP1 constructs (**Fig. 2A**). In an ERE assay, *ESR1* fusions (ESR1-DAB2, ESR1-SOX9, and ESR1-YAP1) demonstrated significantly enhanced ligand-independent activity after the reduction of endogenous ER (displayed results are relative to ESR1-WT cells treated with siESR1) (**Fig. 2B**). Knockdown of endogenous ER in MDA-MB-134-IV revealed a ligand independent, endocrine resistant, hyperactive phenotype in the ESR1-SOX9 and ESR1-YAP1, but not the ESR1-DAB2 transient cell lines (**Figs. 2C-D**).

**Figure 2:**
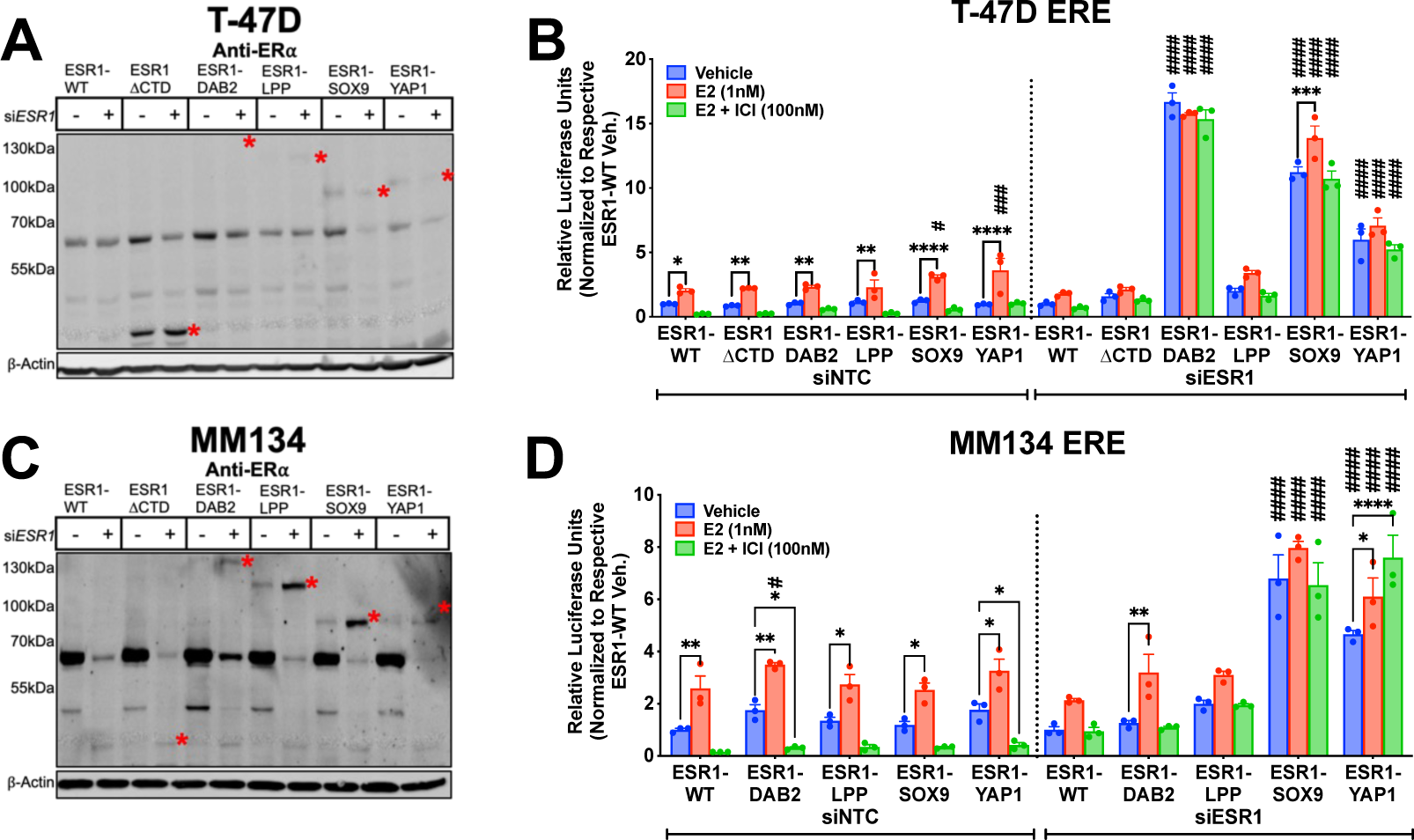
Increased fusion hyperactivity in T-47D and MDA-MB-134-IV cells deplete of endogenous ER. (A) Anti-ER immunoblot of transient ESR1 siRNA or scramble siRNA transfection in T-47D cells stably overexpressing ER constructs. (B) ERE assay in T-47D cells stably expressing ER constructs transfected with non-targeting control (NTC) siRNA or siRNA targeting endogenous ER (siESR1). Raw luminescence values normalized to internal control, renilla luminescence, and to the representative siNTC or siESR1 transfected ESR1-WT vehicle condition. Representative experiment shown with bar graph equivalent to readings ± SEM, n=2. (C) Anti-ER immunoblot of transient ESR1 siRNA or scramble siRNA transfection in MDA-MB-134-iV (MM134) cells stably overexpressing ER constructs. Construct proteins denoted by red asterisks in all blots. β-actin serves as loading control in all blots. (D) ERE assay in MM134 cells stably expressing ER constructs transfected with NTC or siESR1. Normalization and control as above. Representative experiment shown with bar graph equivalent to readings ± SEM, n=2.2way ANOVA with posthoc Dunnett’s multiple comparisons test showing enrichment in comparison to corresponding ESR1-WT treatment (#) or to intra-construct vehicle treatment (*) in each assay. */#p=0.0332, **/##p=0.0021, ***/###p=0.0002, ****/####p<0.0001.

### Model development of stably expressed ER fusions in T-47D and MDA-MB-134-IV cell lines with endogenous ER knockdown

We next stably expressed ER fusions and controls through lentiviral infection of T-47D and MDA-MB-134-IV cell lines. To eliminate the interference of endogenous ER, we reduced endogenous ER levels using short hairpin RNA (shRNA) targeting the 3’ untranslated region of *ESR1* similar to the transient siRNA. Knockdown of endogenous ER in NST T-47D cells correlated with increased levels of *ESR1* fusion proteins (**Fig. 3A**), potentially secondary to increased cellular machinery availability for fusion stability. Treatment with 100nM fulvestrant in the stable T-47D cell lines degraded residual endogenous ER protein expression and ESR1-WT but had little to no effect on ER fusion levels (**Fig. 3B**), indicating that activity of ER fusions exposed to E2, or endocrine therapies, is not confounded by changes in protein level. Compared to the T-47D shESR1 cell lines, MDA-MB-134-IV infected shESR1 cells did not have a corresponding increase in ER fusion expression (**Fig. 3C**).

**Figure 3:**
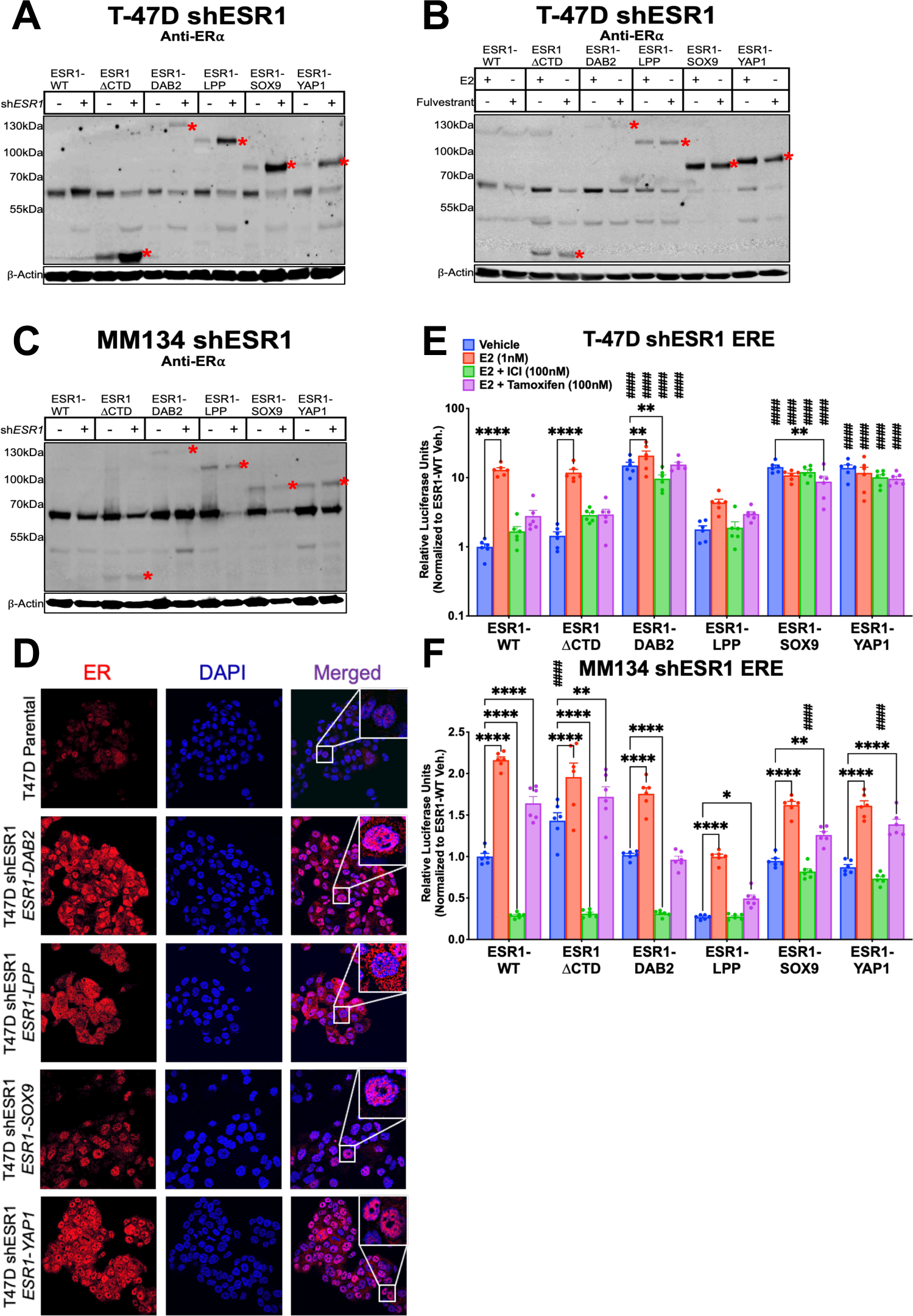
Stable ER fusion overexpression in T-47D and MDA-MB-134-IV endogenous ER depleted cellular contexts. (A) Anti-ER immunoblot of stably overexpressed ER constructs in T-47D cell line stably infected with shESR1. (B) Anti-ER immunoblot of stable T-47D shESR1 cells treated in the presence of 1nM E2 or 100nM fulvestrant. (C) Anti-ER immunoblot of stably overexpressed ER constructs in MM134 cell line stably infected with shESR1. Construct proteins denoted by red asterisks in all blots. β-actin serves as loading control in all blots. (D) T-47D parental and stable shESR1 cells stabling expressing plasmid constructs stained with anti-ER (red) and DAPI (blue) and imaged by confocal microscopy with a 40X objective. Merge represents overlay of anti-ER and DAPI staining. Representative experimental images, n=2. (E) ERE assays in stable T-47D shESER1 cells and (F) MM134 shESR1 cells expressing control and fusion constructs. Raw luminescence values normalized to internal control renilla luminescence and to the corresponding ESR1-WT vehicle condition luminescence readings. Representative experiment shown with bar graph equivalent to readings ± SEM, n=3 for each experiment. 2way ANOVA with posthoc Dunnett’s multiple comparisons test showing enrichment in comparison to corresponding ESR1-WT treatment (#) or to intra-construct vehicle treatment (*) in each assay. */#p=0.0332, **/##p=0.0021, ***/###p=0.0002, ****/####p<0.0001.

Before studying phenotypes, we performed immunofluorescence to examine localization of ER fusions with parental T-47D cell lines as a control for nuclear endogenous ER (**Fig. 3D**). ESR1-DAB2, ESR1-SOX9 and ESR1-YAP1 demonstrated nuclear localization whereas ESR1-LPP revealed both nuclear and cytoplasmic ER staining (**Fig. 3D**). ESR1-DAB2, ESR1-SOX9 and ESR1-YAP1 demonstrated robust ligand independent ERE hyperactivity in the NST cell line T-47D shESR1 (**Fig. 3E**), whereas in MDA-MB-134-IV shESR1, hyperactivity was only observed for ESR1-SOX9 and ESR1-YAP1 (**Fig. 3F**). MDA-MB-134-IV shESR1 ESR1-DAB2 ERE activity did not reveal a ligand independent or endocrine resistant phenotype that was observed in the T-47D shESR1 cells, mirroring the results from transient ER knockdown (**Fig. 2**).

### ER fusion transcriptomic profiling reveals unique and shared estrogen response signatures in T-47D shESR1 cells

To assess unique and shared activities of each fusion, RNA sequencing (RNAseq) was performed on the T-47D shESR1 cell lines treated with vehicle or 100nM fulvestrant which allowed us to assess any potential remaining endogenous ER activity and endocrine therapy effects on ESR1-WT expressing cells. Clustering of the top 500 differentially expressed genes (DEGs) revealed three primary groupings: 1) ESR1-WT, ESR1-ΔCTD and ESR1-DAB2, 2) ESR1-LPP and ESR1-YAP1 and lastly 3) ESR1-SOX9 (**Fig. 4A**). ESR1-WT subclustered independently from ESR1-ΔCTD, indicating an effect of the 3’ C-terminal deletion on transcriptional activity. Each construct clustered together irrespective to treatment condition. This was further confirmed by principal component analysis (**Fig. 4B**), suggesting that fulvestrant had a minimal effect on transcriptional activity and that there was minimal response from endogenous ER. Thus, for subsequent analyses to determine DEG and pathway enrichment, the treatment groups (vehicle and fulvestrant) in each cell line were combined as one entity. There was no significant difference in RNAseq denoted *ESR1* expression between cell lines expressing the different *ESR1* constructs (**Supplementary Fig. S2**). ESR1-SOX9 and ESR1-YAP shared the most upregulated DEGs as predicted by global unbiased hierarchical clustering. A total of 463 shared DEGs were enriched in the four ER fusions cell lines compared to both ESR1-ΔCTD and ESR1-WT (**Fig. 4C**).

**Figure 4:**
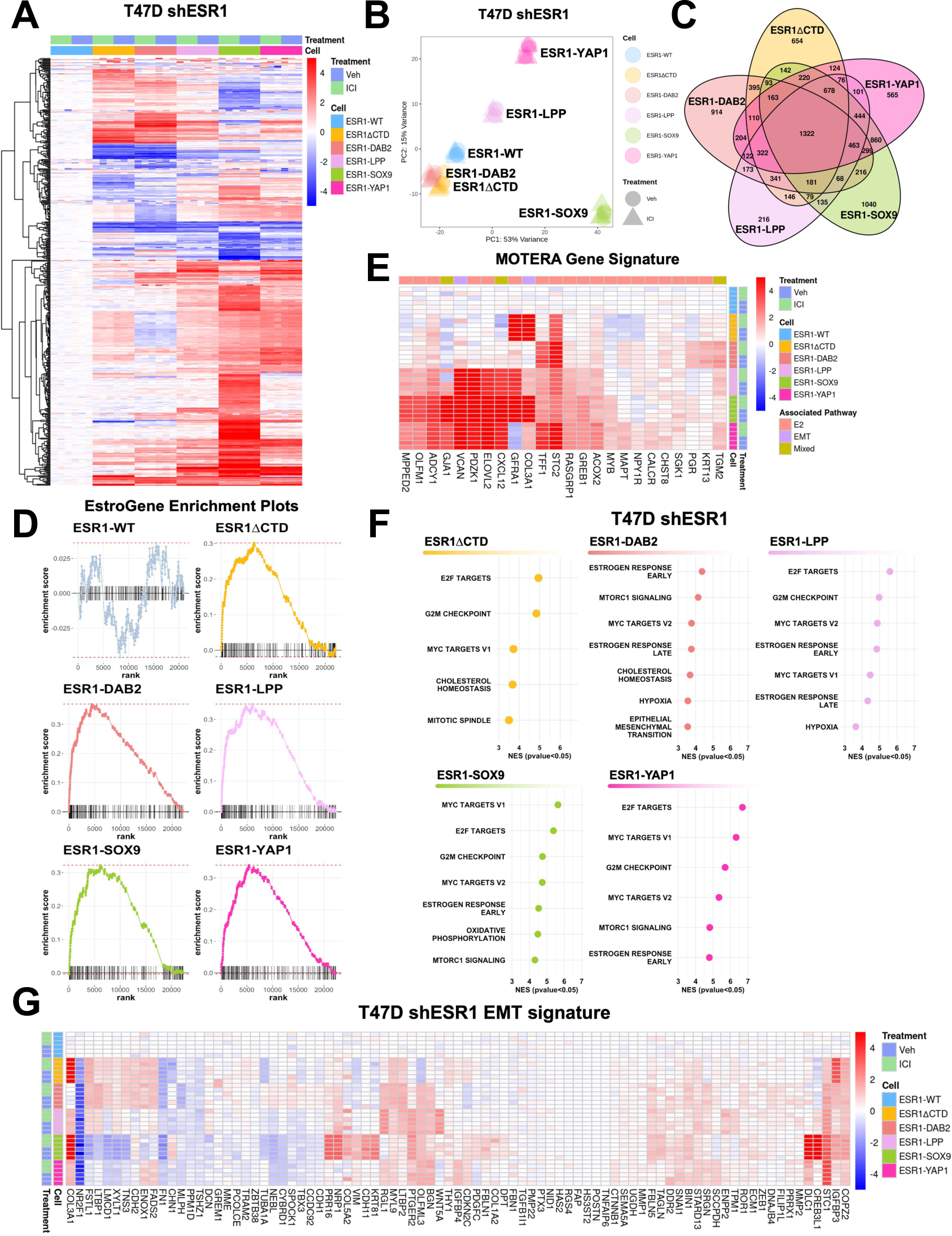
Transcriptomic profiles of ER fusions in ER depleted T-47D cell line enriched in estrogen driven pathways. (A) Heatmap of the top 500 differentially expressed genes (DEG) between vehicle and 100nM ICI treated cells with biased hierarchical clustering of columns. Data normalized to ESR1-WT for relative up- and down-regulation. (B) Principal component analysis (PCA) of T-47D shESR1 ER constructs in the presence or absence of 100nM ICI treatment. (C) Venn diagram of ER construct DEGs. DEGs generated between ESR1-WT and each ER construct (p-value < 0.05, FC > 0). (D) Enrichment plots for ER constructs using the EstroGene signature. ESR1-WT DEGs between treatment groups used as negative control. (E) Heatmap of the 24 gene signature, MOTERA. Data normalized to ESR1-WT for relative up- and down-regulation. (F) CSEA analysis dotplots of fusion cell lines utilizing the DEGs between each construct and ESR1-WT. Size of dot has no significance. X-axis denotes normalized gene enrichment scores (NES). All pathways plotted have NES p-value < 0.05. DEGs generated between ESR1-WT and each ER construct (p-value < 0.05, FC > 0). (G) Heatmap of EMT related genes. Data normalized to ESR1-WT for relative up- and down-regulation.

To assess how ER fusions affect the transcriptome, we compared DEGs to previously reported estrogen response signatures. Our laboratory has recently established EstroGene, a database containing publicly available RNA sequencing, ChIP sequencing and Assay for Transposase-Accessible Chromatin (ATAC) sequencing data in breast cancer cell lines treated with vehicle control or estradiol at various dosages and durations (28). From this dataset, the top 10 percentile of estradiol regulated genes in T-47D cells (n=127) was utilized as an estrogen response signature.

DEGs from ER fusions were significantly enriched in the EstroGene pathway signature in comparison to ESR1-WT DEGs when performing enrichment analysis with the IndepthPathway module’s calculated gene weight values (**Fig. 4D**) (29). Notably, ESR1-WT was not enriched as expected given the absence of E2 and treatment with fulvestrant in the experiment. We also sought to examine the signature Gou et. al. developed to determine ER fusion activity, MOTERA (Mutant or Translocated Estrogen Receptor Alpha) (26). Supervised clustering revealed the enrichment of MOTERA in each of the ER fusions in comparison to ESR1-WT (**Fig. 4E**).

To better understand the contribution of the ER fusion partner to the chimeric protein function, we examined the ER fusion DEGs in a variation of gene set enrichment analysis (GSEA) that again utilized the IndepthPathway module algorithm (29). In brief, the statistical analyses from each inputted DEG list were used to develop a weighted gene list and to perform subsequent concept signature enrichment analysis (CSEA) using a collection of pathway gene sets (29). The top 50 pathways up- or down-regulated are disambiguated by assessing pathway crosstalk and mediating the degree of overlap in numerous pathway gene sets. Disambiguation analyses used a p-value of less than 0.01 (29). The resulting CSEA list is a significant and precise collection of pathways enriched in each individual fusion construct compared to ESR1-WT.

Pathways presented are from the MSigDB Hallmark collection for simplification as these seemed most applicable for interpretation into fusion functionality.

ESR1-DAB2, ESR1-SOX9 and ESR1-YAP1 harbored significant enrichment in mTOR signaling and in all fusion constructs, the estrogen response pathway early was significantly enriched compared to ESR1-WT (**Fig. 4F**). The oncogenic MYC pathway was upregulated in all the ligand depleted transcriptomes of the ER fusions depleted of LBD (including ESR1-ΔCTD). Both ESR1-DAB2 and ESR1-LPP DEGs were enriched in hypoxia pathways with normalized enrichment scores (NES) of 3.54 and 3.63, respectively. The hallmark EMT signature was only enriched in the disambiguated CSEA lists derived from the ESR1-DAB2 DEG set (NES value of 3.53). EMT was significantly enriched by CSEA in ESR1-LPP, ESR1-SOX9 and ESR1-YAP1 pathway analysis, but not enriched with further disambiguation (NES values of 3.62, 2.93 and 2.46 respectively). Although the Hallmark EMT pathway was not enriched after disambiguation analysis in all fusions, genes contributing to the EMT signature were largely upregulated in fusions compared to ESR1-WT (**Fig. 4G**). Importantly, *SNAI1* was upregulated both at the RNA and protein level consistent with literature (**Fig. 4G, Supplementary Fig. S3**) (26). The overall similarity of these independent pathway enrichments derived from unique transcriptomes raises promise regarding potential shared drug susceptible among fusion proteins irrespective of the 3’ partner gene.

Gap junction *GJA1* was also found to be enriched in fusion positive cells previously described by the Ellis group (26). Recent work by Li et. al. uncovered an important role of gap junction proteins in *ESR1* point mutation (Y537S and D538G) driven tumor progression (14). ESR1-SOX9 cells were markedly enriched in the connexin gene family, suggesting a unique functionality to ESR1-SOX9 that remains to be further investigated (**Supplementary Fig. S4**). In summary, transcriptomic profiling of T-47D cells overexpressing ER fusion proteins demonstrated pathway enrichment shared between the fusions, as well as unique changes which might contribute to the varying levels of endocrine resistant activity.

### Cistromic profiling reveals unique transcriptional regulation by ER fusions compared to ESR1-WT

To better understand ER fusions activity as transcription factors, chromatin immunoprecipitation sequencing (ChIP-seq) was performed using the HA tag present in all ER constructs. Hormone deprived ESR1-WT cells treated with 1nM E2 for 45 minutes served as a control for the E2 induced ER cistrome; all other cells were examined under vehicle conditions i.e., without E2. All ER fusions displayed enriched binding at the well-known E2 regulated ER binding sites in *GREB1* and *IGFBP4* (**Supplementary Fig. S5**). Furthermore, ligand depleted ESR1-LPP, ESR1-SOX9 and ESR1-YAP1 had enhanced ChIP-seq signals at ER binding sites, at an intensity similar to that of the estradiol treated ESR1-WT cell cistrome (**Fig. 5A**). To further investigate fusion specific cistromes, the overlap of canonical ER binding sites was identified between each fusion protein and ESR1-WT. As expected, ESR1-WT treated with estradiol resulted in an enhanced number of unique ER binding regions compared to vehicle treated ESR1-WT cells (**Fig. 5B**). Both ESR1-ΔCTD and ESR1-DAB2 shared more ER binding regions with ESR1-WT compared to new unique binding sites. ESR1-LPP shared 105 binding regions with ESR1-WT, but also was bound to 117 unique binding regions. Surprisingly, ESR1-SOX9 had a strong overlap with ESR1-WT (57.5%) and relatively few unique ER binding sites, however, of these binding sites, 94% contained ERE motifs (**Supplementary Table S1**). ESR1-YAP1 had 1,040 unique ER binding sites compared to ESR1-WT; 23% of the unique binding regions harbored ERE motifs suggesting a more promiscuous binding to ER associated regions irrespective of ERE.

**Figure 5:**
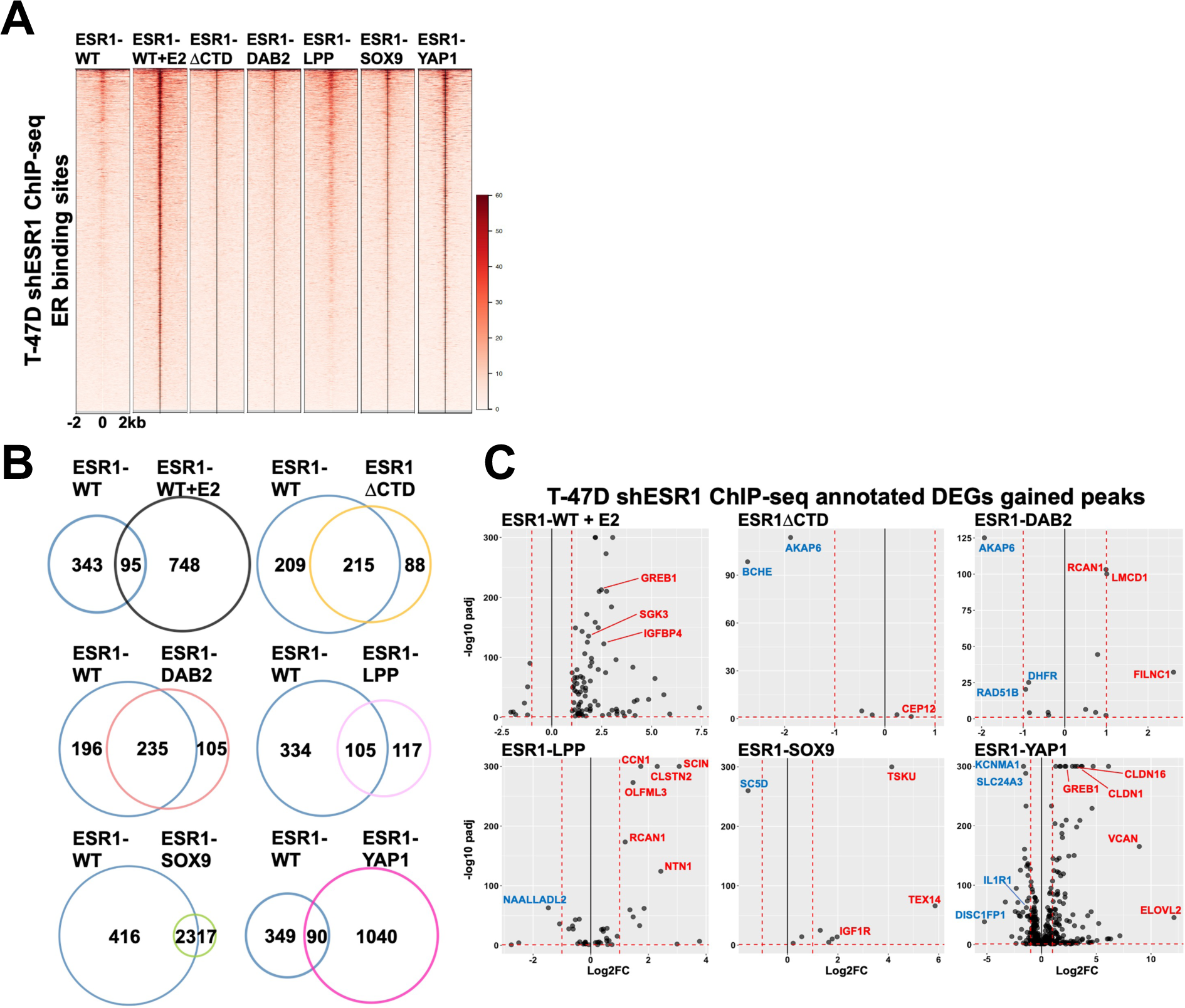
Fusion cistromes reveal direct fusion transcriptomic functionality. (A) Heatmap of ER binding intensities on T-47D shESR1 HA ChIP-seq detected sites in ESR1-WT and ER constructs. Displayed in horizontal window of ±2kb from peak center. (B) Venn diagrams displaying overlap of shared and unique genomic binding sites between ESR1-WT and ESR1-WT stimulated with 1nM E2 or ESR1-WT and ER constructs. (C) Volcano plots of genes up- and down-regulated in both transcriptomic and cistromic analysis. Select up-regulated genes highlighted in red and select down-regulated genes highlighted in blue.

The unique binding peaks of each *ESR1* fusion compared to ESR1-WT were then integrated with their associative transcriptomes in order to develop a list of DEGs that could help explain fusion transcriptional regulation at gene loci. The ESR1-DAB2 cistromic-transcriptomic overlap revealed enrichment of tumor suppressor gene, *RCAN1*, as well as an upstream negative regulator of Myc-1, *FILNC1* (**Fig. 5C**) (30). ESR1-LPP was also found to have an enhanced binding and subsequent transcription of *RCAN1*. In addition, however, ESR1-LPP was enriched in transcription of oncogenic *CCN1* and *NTN1*. ESR1-SOX9 cistromic-transcriptome revealed downregulation of a *SC5D* which is a positive prognostic indicator of neoadjuvant chemotherapy response (31). The cistromic-transcriptomic overlap also revealed enrichment of *IGF1R* in ESR1-SOX9 expressing cells. Our research group has studied the role of *IGF1R* in the context of different molecular subtypes of breast cancer (32–36). The enrichment of the gene provides a potential novel axis to target ESR1-SOX9 positive tumors. The fusion which had the most congruence between cistromic and transcriptomic analyses was ESR1-YAP1, suggesting direct transcriptional activity. Analysis revealed an upregulation of a chemoresistant biomarker, *CLDN16* (37). In addition, another claudin family member, *CLDN1*, was upregulated in ESR1-YAP1 cells. *CLDN1* has been described as downregulated in luminal breast cancer, but upregulated in aggressive basal like subtypes which may potentially suggest subtype switching for cells expressing ESR1-YAP1 or other ER fusions (38). ESR1-YAP1 cistromic-transcriptomic overlap also revealed enrichment of *ELOVL2,* a tumor suppressor in tamoxifen resistant settings, which emphasizes the need for further investigation in how tumor suppressors may be functioning in fusion dependent contexts (39). ESR1-YAP1 was found to have enhanced peak enrichment at the *GREB1* and *VCAN* locus supporting the fusion’s functionality in estrogen mediated and migratory pathways. Given the similarity in pathway enrichment between the fusion constructs, the differences in fusion cistromes implies a diversity in upstream genomic regulation that ultimately results in shared cellular activity.

### The contribution of ER fusion proteins to oncogenic phenotypes are assay dependent and cell line specific

To understand the contribution of ER fusions to endocrine resistance and metastatic phenotypes, we examined growth, cell survival, and cell migration in the NST and ILC stable cell line models. Consistent with ERE hyperactivity, T-47D shESR1 ESR1-SOX9 and ESR1-YAP1 cell lines displayed greater proliferation compared to ESR1-WT, irrespective of treatment condition (**Fig. 6A**). ESR1-LPP had a more consistent ligand independent, endocrine resistant, growth pattern in both 2D and 3D conditions compared to ESR1-DAB2 (**Fig. 6A**). Consistent with the growth phenotype, T-47D shESR1 ESR1-LPP, ESR1-SOX9, and ESR1-YAP1 stable cell lines displayed enhanced colony formation in hormone deprived conditions and at low cellular densities compared to ESR1-WT (**Fig. 6B**). ESR1-DAB2, analogous to the growth assay, did not have a robust colony formation phenotype (**Fig. 6B**). We next assessed migration of the T-47D shESR1 cells employing a wound scratch assay. Although not significant, at the 72-hour timepoint post scratch creation, there was a consistent trend in migratory closure in the ESR1-SOX9 and ESR1-YAP1 cell line models (**Fig. 6C**). Both ESR1-DAB2 and ESR1-LPP cell migration was similar to that of the ESR1-WT cell line.

**Figure 6:**
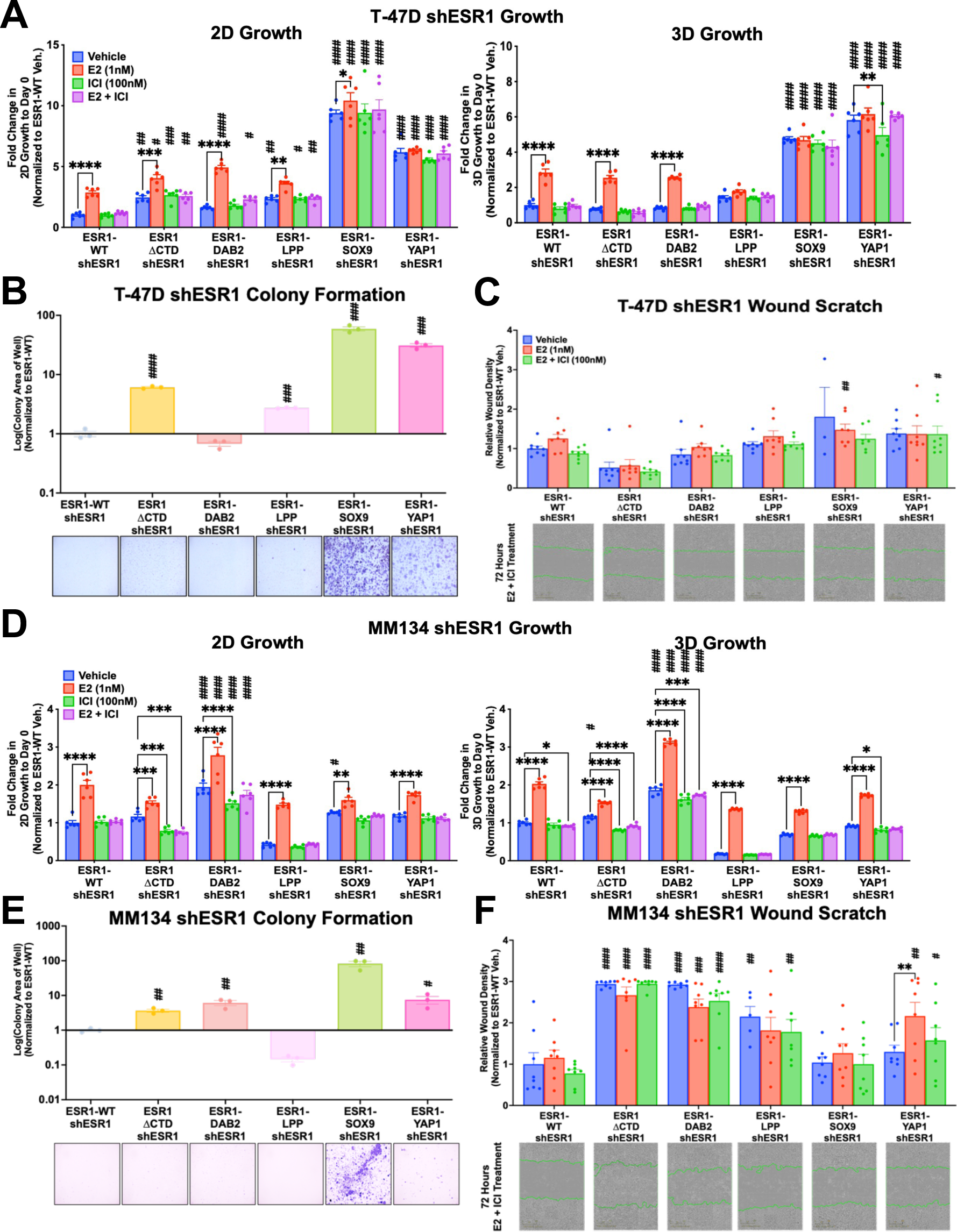
Cellular context dependent fusion enrichment in metastatic properties. 2D and 3D growth in T-47D (A) and MM134 (D) shESR1 cell lines. Cell viability was quantified on days 0 and 9 (T-47D) or 4 (MM134) with CellTiter-Glo assay and data was normalized to day 0 quantification. Representative experiment shown, n=3 (each with six technical replicates). (B) T-47D shESR1 and (E) MM134 shESR1 colony formation assays. Cells were seeded at a low cell density and stained with 0.5% Crystal Violet at approximately 2 weeks (T-47D) or 3.5 weeks (MM134). Representative images are shown, n=3 (each with three technical replicates). Wound scratch healing assay of (C) T-47D shESR1 and (F) MM134 shESR1 cells in the presence of anti-proliferation treatment with mitomycin C. Bar graphs generated from IncuCyte automated program to calculate relative wound density after a 72-hour time period. Representative images from the E2 and ICI treated cohort. Green outline represents initial wound scratch leading edges, n=3 (each with eight technical replicates). 2way ANOVA with posthoc Dunnett’s multiple comparisons test showing enrichment in comparison to corresponding ESR1-WT treatment (#) or to intra-construct vehicle treatment (*) in each assay. */#p=0.0332, **/##p=0.0021, ***/###p=0.0002, ****/####p<0.0001.

To also characterize the fusions in an ILC cell line, we performed the same assays in the MDA-MB-134-IV shESR1 cell lines stably expressing the ER constructs. Endocrine resistant growth was not evident in the MDA-MB-134-IV shESR1 cells, except for ESR1-DAB2 in comparison to ESR1-WT (**Fig. 6D**). Although ESR1-SOX9 and ESR1-YAP1 did not display enhanced growth phenotypes, the ER fusions did demonstrate significantly enhanced colony formation compared to ESR1-WT (**Fig. 6E**). We also observed more robust colony formation in ESR1-DAB2 cells suggesting a greater affinity to cluster together. This in part could be explained by a significantly enhanced migratory phenotype of the ESR1-DAB2 MDA-MB-134-IV shESR1 cells (**Fig. 6F**). Interestingly, ESR1-ΔCTD in the MDA-MB-134-IV shESR1 cells also presented a higher migratory phenotype then ESR1-WT (**Fig. 6F**). The stark difference between the NST T-47D and ILC MDA-MB-134-IV fusion cell lines requires further investigation in the cellular mechanisms of promoting unique *ESR1* fusion protein functionalities (**Fig. 7**).

**Figure 7:**
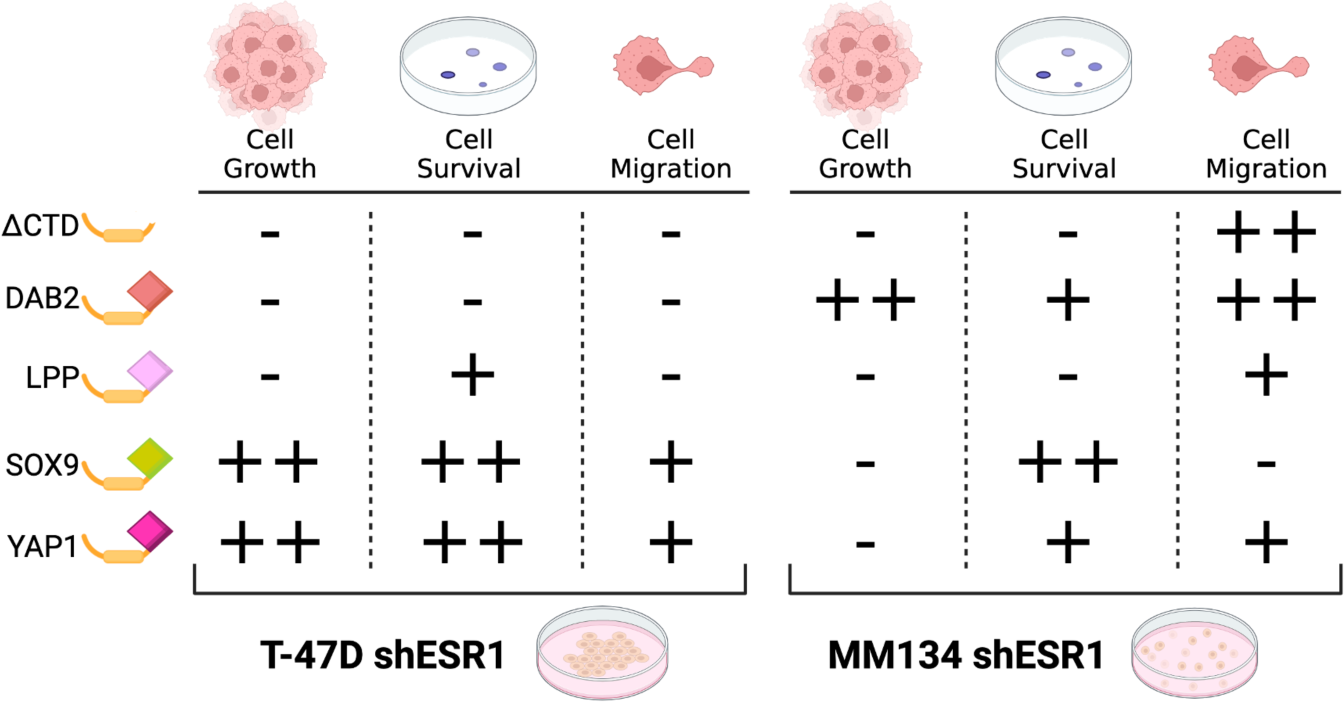
ER fusion metastatic phenotype summary. Schematic of ER fusions and corresponding oncogenic activity in T-47D and MM134 cell lines. Plus sign indicates activity. Minus sign signifies no appreciable activity. Activity is estimated in comparison to ESR1-WT in ligand independent and endocrine resistant conditions.

## Discussion

Endocrine resistance develops in a third of patients diagnosed with primary ER positive disease and furthermore, in the majority of patients with metastatic disease (40). While ER point mutations respond to estrogen and anti-estrogens, ER fusion proteins lack a hormone binding domain and thus represent a mechanism of pure ligand independence and endocrine resistance. Herein, we report that four ER fusion proteins are functionally active in HEK293 cells irrespective of estradiol or endocrine therapy stimulation. These results held true in C4-12 cells that are devoid of endogenous ER. In NST T-47D and ILC MDA-MB-134-IV cell line models, transient and stable knockdown of endogenous ER enabled dissection of fusion specific phenotypes. The robust ERE response pattern in T-47D shESR1 cells expressing ESR1-DAB2, ESR1-SOX9 or ESR1-YAP1 differed from equivocal ILC cell lines suggesting a molecular subtype influence on fusion function. Through RNA sequencing, we report that T-47D shESR1 fusion transcriptomes subcluster from ESR1-WT in estradiol depleted or ICI treated conditions. The transcriptionally active fusions in the ERE assays, ESR1-SOX9 and ESR1-YAP1, demonstrated the greatest overlap in differentially expressed genes. All fusions shared estrogen and oncogenic pathway enrichment. The direct and indirect capacity of fusion activity varied, however, unsurprisingly, fusion partners that were transcription factors harbored the most direct DNA binding activity. Lastly, we report on the metastatic phenotypes of fusions in the NST and ILC shESR1 cell line models. These assays provided the greatest insight to differences in a fusion’s function depending upon histological subtype and phenotype being assessed. ILC is the second most prevalent histologically subtype of breast cancer and is predominantly ER positive. ER point mutations developed in metastatic disease do not differ in type nor prevalence between NST and ILC tumors; therefore it is imperative to study fusions in both representative disease models (41). ILC tumors have higher extracellular stromal deposition, are distinguished by loss of tight junction E-cadherin, and are enriched in HER2, HER3, FOXA1, and PIK3CA mutations as well as PTEN loss (42). The MDA-MB-134-IV cell line furthermore harbors KRAS and MAP2K4 mutations. We and colleagues reported an increased sensitivity to chaperone heat shock protein size 70 kDa (HSP70) inhibition in MDA-MB-134-IV cells comparative to T-47D cells, suggesting a potential mechanism of fusion or downstream transcriptome stabilization, particularly in a setting of elevated proteomic stress (43). Our present work affirms that the distinct molecular landscape of an ILC cell line influences fusion behavior. ESR1-LPP demonstrated an ILC specific migratory phenotype in comparison to lack of activity in the NST cell line model. Similarly, a reproducible significant growth enrichment was appreciated in ILC ESR1-DAB2 cells. Gou et. al. described differences in ERE activity of ESR1-DAB2 between two NST cell lines but, ultimately described the fusion as inactive given transcriptomic data and their MOTERA signature (26). Our independent, pre-clinical experimentation in an ILC model, albeit in a single cell line, provides the sole insight into fusion mechanistic variation between cancer molecular landscapes. Further investigation into subtype specific *ESR1* chimeric protein functionality in both *in vitro* and *in vivo* models is warranted. Since the discovery of an ESR1-YAP1 fusion in 2013 (25), numerous *ESR1* exon 6 fusion events have been identified, however, few have been comprehensively studied. Lei et. al. reported increased transcriptional activity of fusions ESR1-YAP1, ESR1-PCDH11X and ESR1-NOP2 (44). A co-immunoprecipitation assay revealed a lack of endogenous ER-fusion dimerization, however a biotinylated ERE pulldown confirmed ER fusion binding at EREs (44). RNAseq and confirmatory ChIP-seq identified enrichment in EMT and estrogen response pathways in the two phenotypically active fusions (44). The authors further tested *in vitro* and *in vivo* inhibition of fusion activity with a CDK4/6 checkpoint inhibitor, Palbociclib, which demonstrated dose-dependent growth inhibition and remains to be further evaluated as a therapeutic in other identified *ESR1* fusions (44). The same research group expanded on their findings, again in NST cell lines and patient derived xenograft models, with a focus on a fusion activity transcriptomic signature MOTERA (26). In the MOTERA signature, the majority of genes are estrogen responsive genes in the classical ER signaling network. A few genes, *VCAN* and *COL3A1*, are components of the EMT pathways. Regardless of the lack of dimerization between endogenous ER and *ESR1* fusions, endogenous ER presumably competes for DNA binding sites or squelches ER co-activators, thus complicating interpretation of fusion driven cellular activity. Utilizing cells with depletion of endogenous ER, we found a more robust fusion phenotype that was independent of the endogenous ER influence and that was furthermore breast cancer subtype dependent.

The enrichment of a E2 related signature in Gou et al.’s work compelled us to examine the expression levels of a cohort of up-regulated E2 responsive genes derived from T-47D RNAseq data that is publicly available through EstroGene (28). All four fusion constructs and the truncated fusion protein were enriched in the E2 signature emphasizing that ER fusions retain components of the canonical ER regulome. ESR1-ΔCTD’s EstroGene enrichment is potentially contributable to non-canonical pathways (i.e., through alternatives to direct ERE activation) that result in EstroGene signatures. This is particularly interesting in MDA-MB-134 cells given the colony formation and migration phenotypes of ESR1-ΔCTD. It is possible that if ESR1-ΔCTD binds ERE motifs, the lack of size (i.e., absence of a 3’ fusion partner), may inhibit the truncated protein’s ability to serve as an active transcription mediator. The diverse nature of ER fusion 3’ partners are hypothesized to influence unique fusion-specific transcriptomes. Perhaps unexpected, however, was the similarity in pathway convergence among the hormone deprived fusion transcriptomes, including upregulation of proliferative and EMT related pathways. MYC (v-myc avian myelocytomatosis viral oncogene homolog) targets is a gene set enriched in all ligand depleted *ESR1* chimeric cell lines compared to ESR1-WT, suggesting a potential effect of the 5’ portion of *ESR1* on pathway regulation. The MYC pathway has been implicated in breast cancer progression and is associated with poor prognosis (45). Furthermore, MYC signaling has been described in anti-estrogen resistance and distant relapse of ER positive disease (46). The consistency in the upregulation of the MYC pathway, even after disambiguation, shows a promising overlap between the otherwise distinct phenotypic responses from each fusion. ESR1-DAB2 was the most distinct fusion in terms of unique pathway enrichment such as enrichment in hypoxia and cholesterol homeostasis, consistent with previously reported RNAseq results (26). The enrichment of these metastatic potentiating pathways, independent of the estrogen signaling network, emphasizes the importance of a multifaceted experimental approach to determine fusion functionality holistically.

The transcriptome of ESR1-LPP was surprisingly more similar to ESR1-SOX9 and ESR1-YAP1 than hypothesized based upon phenotypic assays in which ESR1-LPP driven activity was comparatively reduced. ESR1-LPP was found to be both nuclear and cytoplasmic. Thus, the fusion may be involved in regulating extra-nuclear functions that ultimately converge on similar global transcriptomic profiling as ESR1-SOX9 and ESR1-YAP1. Another unexpected finding was the enrichment of collagen type III alpha 1 chain (*COL3A1*) in ESR1-ΔCTD positive cells. *COL3A1* is a member of the EMT regulome that is associated with poor prognosis in ER positive breast cancer (47). In T-47D cells, ESR1-ΔCTD displayed no enhanced migration, however, in MDA-MB-134-IV cells, there was significantly elevated motility. Whether *COL3A1* influences an enhanced MDA-MB-134-IV ESR1-ΔCTD phenotypic response requires further experimentation. ESR1-SOX9 was additionally enriched in *COL3A1* expression as well oxidative phosphorylation and cell adhesion pathway members. We have previously published on enhanced cell-cell adhesion networks in *ESR1* point mutant models that translated to increased circulating tumor cell clusters in patients with metastatic, endocrine resistant disease (14). The similarity of heightened cell adhesion properties in *ESR1* fusions supports a potential avenue to targeting all functioning *ESR1* chimeras with disruption of the LBD. Finally, Gou et. al. demonstrated an enrichment in ERE motifs in their ESR1-YAP1 T-47D model ChIP-seq, consistent with our analysis (26).

In the present study, beyond the consistent transcriptomic profiling outlined above, we demonstrated congruent functional phenotypes to that of Gou et. al. and Lei et. al. (26, 44). Growth, cell survival, and migration in the ESR1-SOX9 and ESR1-YAP1 T-47D shESR1 cell lines was consistent with the *in vitro* transcriptional activity previously published for NST models; these fusions were consistently the most active. Importantly, our work is compared to a stably expressed ESR1-WT plasmid in T-47D or MDA-MB-134-IV shESR1 cells rather than using a YFP control. As previously described, ESR1-DAB2 displayed context dependent activity. DAB2 positive tumor-associated macrophages (TAMs) are associated with overall poor clinical outcome in ILC (48). The DAB2 canonical function in the TAMs leads to extracellular matrix (ECM) remodeling in a pro-metastatic fashion (48). Although this phenomenon has been strictly reported in TAMs, there is the potential that the DAB2 fusion protein remodels the ECM in the higher stromal microenvironment of ILC cell lines. Consistent with this notion is ESR1-DAB2 cell line enrichment in EMT, which may enable enhanced phenotypes in the MDA-MB-134-IV cellular context. Surprisingly, ESR1-SOX9 did not have as robust phenotypes in growth and migration in the ILC MDA-MB-134-IV cells. Interestingly, cell survival was enriched in all T-47D constructs besides ESR1-DAB2, which may be contributed to the lack of upregulation in E2F target genes.

The role of the 3’ partner in fusion functionality remains an active research endeavor. Despite the enrichment of E2 signatures, the diverse functional profiles suggest a predominate influence from the translocated gene. In addition, the fusion models’ enrichment in MTOR pathways was not exhibited in ESR1-ΔCTD suggesting a functional or stabilizing role of the 3’ partner, whereas MYC related pathways may be contributed to *ESR1* exons 1-6. Annotated cistrome DEGs revealed ESR1-YAP1 regulated transcription of the canonical YAP1 regulome including fatty acid elongation and claudin family members, indicating a canonical influence of the YAP1 transcription factor on the fusion transcriptional activity. This may help explain the enhanced functional activity that we and others have reported in the ESR1-YAP1 model. Similarly, enhanced ESR1-SOX9 activity may be secondary to the 3’ partner’s role as a transcription factor, however, potentially through a co-activating or stabilizing function as the ESR1-SOX9 cistrome was enriched in ERE binding motifs. The selective affinity of ESR1-SOX9 for binding EREs in an E2 depleted context suggests a narrow, yet influential, regulation of ER related gene regions.

Together, our results augment the limited literature on ER fusion proteins in advanced treatment resistant breast cancer. Critically, our work emphasizes the uncovered role in which cellular context and phenotypic properties contribute to observed ER fusion action. Continued investigation of ER fusion proteins in both NST and ILC contexts may provide alternate treatment regimens in patients diagnosed with endocrine resistant breast cancer; an acutely relevant endeavor in the setting of updated guidelines to identify ER mutants during therapeutic intervention (49).

## Methods

### Cell culture

Cell lines employed in our research are sourced from American Type Culture Collection (ATCC): HEK293T (RRID: CVCL_0045), T-47D (RRID: CVCL_0553) and MDA-MB-134-VI (RRID: CVCL_0617). All cells were maintained in media supplemented with 10% fetal bovine serum (FBS; Life Technologies #26140079): HEK293 in Dulbecco’s modified Eagle Medium (DMEM, Life Technologies #11965118); T-47D in Roswell Park Memorial Institute (RPMI, Life Technologies #11875-119) 1640 and MDA-MB-134-IV in 50:50 composition of DMEM and Leibovitz’s L-15 medium (Invitrogen #11415-114). MCF-7/C4-12 (RRID: CVCL_GX99), is an estrogen-independent MCF-7 (RRID: CVCL_0031) derivative generated by nine months of continuous hormone-deprivation (27). C4-12 cells were cultured in phenol free minimum essential medium (MEM, Fisher #A10488-01) alpha media supplemented with 10% charcoal stripped serum (CSS; Sigma-Aldrich #F6765). Cell lines are maintained at 37°C and 5% CO2. Cells are routinely tested to be mycoplasma free (MycoAlert Mycoplasma Detection Kit (Lonza, Basel, Switzerland # LT07-418)) and authenticated by the University of Arizona Genetics Core (Tucson, Arizona). Hormone deprivation (media change to IMEM supplemented with 10% CSS thrice per day for three days) was performed for all experiments unless otherwise stated.

### Reagents

17*β*-estadiol (E2), purchased from Sigma (#E8875), and Fulvestrant (ICI182,780) obtained from Tocris (#1047) were both reconstituted in 100% ethanol. Tamoxifen (4-hydroxytamoxifen, 4-OHT) was obtained from Sigma (#H6278) and was reconstituted in dimethyl sulfoxide (DMSO; ATCC).

### Plasmid Construction and Stable Cell Line Generation

ER fusion plasmid constructs were engineered by Gene Universal (Newark, Delaware) by custom gene synthesis. Identified *ESR1* fusion sequences were supplied for integration into a pCDH-MSCV-MCS-EF1*α*-copGFP-T2A-Puro backbone vector obtained from Systems Biosciences (#CD713B). The *ESR1* fusion sequence was inserted at NheI and SwaI restriction sites with an upstream Kozak sequence and Human influenza hemagglutinin (HA) tag. A second ESR1-DAB2 plasmid in a pCDH-CMV-MCS-ER1*α*-Puro (System Biosciences #CD510B-1) was received as a kind gift from Dr. Matthew Ellis (AstraZeneca; formal principal investigator at the Baylor College of Medicine, Texas). Lentiviral infection was employed for stable expression of ER chimeric plasmids in T-47D, MDA-MB-134-IV and C4-12 cells. Plasmids were transfected into HEK293T cells with packaging plasmids pMDL, pRev and pVSVG utilizing PEI in a 3:1 PEI to DNA ratio. Incubated target media on packaging cells was collected and filtered through a 0.45µM syringe onto target cell lines at approximately 70% confluency with polybrene (4 µg/ml) at 36 hours, 48 hours, and 72 hours post-transfection. Infected cells were selected with puromycin for at least seven days.

### shESR1 Design and Infection

Short hairpin (sh) RNA targeting the 3’ untranslated region (UTR) of *ESR1* in exon 8 was originally inserted in the MLPE backbone (a kind gift from Scott W. Lowe, Memorial Sloan Kettering Cancer Center, New York): TGCTGTTGACAGTGAGCGACAGCAAGTTGATCTTAGTTAATAGTGAAGCCACAGATGTATTAACTAAGATCAACTTGCTGCTGCCTACTGCCTCGGA. MLPE-shESR1 and the plasmid for subcloning, SGEN (pRRL) (Dr. Johannes Zuber, RRID:Addgene 111171), were digested at XhoI (New England BioLabs Inc. #R0146S) and EcoRI (New England BioLabs Inc. #R3101S) restriction sites. Gel purification using the Monarch DNA Gel Extraction Kit (New England BioLabs Inc. #T1020L) was performed to extract the shRNA sequence and column purification with the QIAprep PCR Purification Kit (Qiagen #28104) was performed to concentrate the digested SGEN plasmid. The purified shESR1 and SGEN DNA were digested with XhoI and EcoRI and subsequently ligated at a molar ratio of approximately 3:1 with 10X T4 DNA Ligase Buffer (New England BioLabs Inc. #B0202S) and T4 DNA Ligase (New England BioLabs Inc. #M020M). Lentiviral infection was employed for stable expression of shESR1 in T-47D and MDA-MB-134-IV cells.

### siRNA Transient Transfection

T-47D and MDA-MB-134-IV cell lines were plated at equal cell densities in 6-well plates (Fisher #08-772-1B) in order to reach 70% confluency in a 24-hour period. Two custom short interfering (si) RNA oligos targeting the 3’ untranslated region (UTR) of *ESR1* were designed and ordered from Horizon Discovery/Dharmacon (Lafayette, Colorado): UCUCUAGCCAGGCACAUUC (sense sequence, ESR1_1) and UCAUCGCAUUCCUUGCAAA (sense sequence, ESR1_2). siGENOME non-targeting siRNA was obtain for a scramble control (Horizon Discovery #D-001206-14). Cells collected for immunoblotting were infected with siRNA for 72 hours before lysis. Cells infected with siRNA for ERE assaying were infected for 72 hours with siRNA before plate reading. Equivalent quantities of siESR1 or scramble siRNA (scRNA) were forward transfected following the Lipofectamine RNAiMAX (ThermoFisher Scientific #13778150) protocol according to manufacturer’s instructions.

### Immunoblotting

Cellular protein lysates were harvested utilizing RIPA buffer (50mM Tris pH 7.4, 150mM NaCl, 1mM EDTA (Thermo Fisher Scientific #15-575-020), 0.5% Nonidet P-40 (Sigma Aldrich #74385), 0.5% sodium deoxycholate, 0.1% SDS) supplemented with 1X HALT protease and phosphatase cocktail (Thermo Fisher #78442). Samples were vortexed, probe sonicated for 15 seconds (20% amplitude, Ultrasonic Processor GEX130) and centrifuged at 14,000rpm at 4°C for 15 minutes. Protein concentration was assessed using the Pierce Bicinchoninic acid (BCA) protein assay (Thermo Fisher #23225). Unless otherwise stated, 50µg of each protein sample was run on a 10% SDS-PAGE gel followed with a 90V transfer at 4°C for 90 minutes to a PVDF membrane (Millipore #IPFL00010). Membranes were blocked for one hour with Intercept PBS blocking buffer (LiCor #927-40000) at room temperature with rocking. Antibody probing was performed overnight at 4°C with rocking: ER*α*, clone 60C (Millipore #04-820, RRID:AB_1587018); HA (C29F4) (Cell Signaling Technologies #3724, RRID:AB_1549585); *β*-actin (Millipore Sigma #A5441, RRID:AB_ 476744); snai1 (I70G2) (Cell Signaling Technologies #3895S). After removal of primary antibodies, blots were wash with 1X PBS-Tween 20 (0.1%) for 15 minutes, three times. Secondary antibodies were applied for a one-hour room temperature incubation (1:10,000; anti-mouse 680LT (LiCor #925-68020); anti-rabbit 800CW (LiCor #925-32211)). Imaging of membranes was performed on the LiCor Odyssey CLx Imaging system.

### ERE-Tk-Luc Assay

C4-12, T-47D, MDA-MB-134-IV cells were seeded in triplicates in 24-well plates (Fisher #08-772-1) at 250,000 cells/well, 220,000 cells/well, and 255,000 cells/well, respectively. HEK293T cells were seeded in replicates of six in 96-well-plates (Falcon #353072) at 37,400 cells/well. Seeding density was calculated in order to reach 70% confluency within 24 hours. 24 hours post plating, cells were washed with 1X PBS and incubated for 24 hours in phenol-red-free modified improved MEM (IMEM, Fisher Scientific #MT10026CV) enriched in 10% CSS for hormone deprivation. Cells were co-transfected at a ratio of 5:1 with a vector harboring a firefly luciferase gene under an estrogen-response-element promoter (ERE-Tk-Luc) and a vector harboring renilla luciferase under a null promoter using the Lipofectamine 3000 transfection method following the manufacturer’s protocol. 24 hours post transfection, cells were treated with either 1nM estradiol (E2), 1nM E2 and 100nM fulvestrant, 1nM E2 and 100nM tamoxifen, or equal volume of 100% ethanol. C4-12, T-47D, MDA-MB-134-IV cells were lysed 48 hours post-transfection with 100µl 1X passive lysis buffer (Promega #E1960) at room temperature for 20 minutes. Following lysis, 20µl of lysed cells were transferred to a white 96 well plate (MIDSCI #781605) in duplicates before measurement using the Dual-Luciferase Reporter Assay (Promega #E1960). HEK293T cells were lysed and assayed using the Dual-Glo kit following the manufacturer’s protocol (Promega #E2940). Luminescence was measured using a two-injector dual system on the GLOMAX-Multi+ Microplate Multimode Reader (Promega). Relative luminescence units were calculated by normalizing oxyluciferin readouts to corresponding renilla measurements and in the HEK293T cells an additional normalization to non-transfected control cells was performed.

### Immunofluorescence staining

T-47D shESR1 cells were plated at a density of 150,000 cells/well on glass coverslips (Fisher #12-545-80P) in 24-well plates. 24 hours post-seeding, cells were fixed on ice with 4% paraformaldehyde (Electron Microscopy Sciences #15710-s) for 30 minutes followed by blocking in blocking buffer (0.3% Triton X-100, 5% BSA, 1X Dulbecco’s PBS (DPBS)) for one hour at room temperature. Primary antibody was incubated overnight at 4°C: ER (1:50 dilution, Lecia Biosystems #NCL-L-ER-6F11, RRID:AB_563706). Secondary antibody incubation was performed for one hour at room temperature: anti-mouse AlexaFluor647 (1:200 dilution, ThermoFisher Scientific #A28181, RRID:AB_2536165). Cells were nuclear stained with Hoechst 33342 (1:10,000 dilution, proxy to DAPI stain, ThermoFisher Scientific #H3570) for 10 minutes. Coverslips were mounted with Aqua-Poly/Mount (Polysciences #18606-20) and slides were imaged on a Nikon A1 confocal microscope with the 40X objective.

### RNA sequencing

Hormone deprived T-47D shESR1 cells were plated at equivalent cell densities in triplicate in 6-well plates in order to reach a 70% confluency at 24 hours. 24 hours post seeding, cells were treated with 100nM of ICI or equivalent volume of 100% ethanol for 24 hours. RNA extraction was performed using the Qiagen RNeasy Mini Kit and RNA concentration was quantified via a Nanodrop 2000 spectrophotometer (Thermo Scientific). Purified RNA samples had library preparation, quality control and sequencing conducted at the UPMC Genome Center (Pittsburgh, Pennsylvania). mRNAseq of paired-end 1×100 base pair reads was employed on a NovaSeq 6000 platform utilizing a flow cell SP-200 to achieve a sequencing depth of 19 million reads/sample. RNA sequencing data is in the process of being deposited in the Gene Expression Omnibus database.

### RNA sequencing analysis

RNA sequencing quality assurance was run using FastQC v0.11.8 (50). Expression quantifications were performed using the Salmon v1.3.0 application (51) with reference index Ensemble GRCh38. RNA gene-level summarizations were generated by tximport v1.16.1 with the reference EnsDb.Hsapiens.v86 (52). Principal component analysis was performed using the variance stabilizing transformation (VST) and plotPCA functions from the R package DESeq2 v1.30.1 (53). VST DESeq2 counts were used to observe unbiased hierarchical clustering using the R function pheatmap v1.0.12 (RRID:SCR_016418). Significantly differential expressed genes (DEGs) were defined as genes with an p-value of less than 0.05. No limitation on fold change (FC) threshold was applied. Venn diagram plots were generated using the VennDiagram package (54). Gene Set Enrichment Analysis (GSEA) was performed using the R package fgsea v1.22.0 (55) with pathway lists derived using the msigdbr package (56) or concept signature enrichment analysis (CSEA) was performed with the IndepthPathway application (29). The IndepthPathway module input was the normalized counts of the DESeq2 object. Data visualization was performed using ggplot2 package (57). EstroGene (estrogene.org) is a curated collection of MicroArray analyses, RNAseq analyses, and ChIP-seq analyses (28).

### Quantitative Real Time Polymerase Chain Reaction (qRT-PCR)

Hormone deprived T-47D shESR1 cells were seeded at 1,000,000 cells/well into 6-well plates. Cells were treated for 24 hours with either 1nM estradiol or an equivalent volume of 100% ethanol. RNA was extracted from cells following the RNeasy Mini Kit (Qiagen #74106) protocol. Complementary DNA (cDNA) was synthesized using the iScript RT Supermix reagent (BioRad #1708890). qRT-PCR reactions were performed utilizing the SybrGreen Supermix (BioRad #1726275). The ΔΔCt method was applied to analyze relative mRNA fold changes to an *RPLP0* internal control.

### Chromatin Immunoprecipitation (ChIP), sequencing, and analysis

T-47D shESR1 cells were plated at 80% to 90% confluency prior to hormone deprivation. Hormone deprived cells were treated with 100nM ICI or equivalent volume of 100% ethanol for 45 minutes. An additional hormone deprived ESR1-WT plate was treated with 1nM E2 for 45 minutes as an experimental positive control. Cells were fixed in 1% formaldehyde (Electron Microscopy Sciences #15710-S) for 10 minutes at room temperature with rocking. The reaction was quenched with 0.125M glycine (Invitrogen #15527-013) for 5 minutes at room temperature with rocking. Post-quenching, cells were washed three times in ice cold 1X DPBS, followed by harvest in 1X DPBS supplemented with protease and phosphatase inhibitors at a 1:100 dilution. Harvested cells were centrifuged at 2,500 rpm for 5 minutes at 4°C. Supernatant was aspirated and cell pellets were resuspended in 1mL Lysis Buffer 1 (50mM Hepes-KOH (pH 7.5), 140mM NaCl, 1mM EDTA, 10% glycerol, 0.5% NP-40/Igepal CA-630, 0.25% Triton X-100) and rotated at 4°C for 10 minutes to enrich for the nuclear fraction (58). Cells were pelleted with a 2,500rpm centrifuge for 5 minutes at 4°C. Pellets were resuspended in Lysis Buffer 2 (10mM Tris-HCL (pH 8.0), 200mM NaCl, 1mM EDTA, 0.5mM EGTA) and rotated for 5 minutes at 4°C (58). Cells were pelleted by centrifugation for 5 minutes at 2,500rpm in 4°C. Supernatant was aspirated and cell pellets were resuspended in 300µl Lysis Buffer 3 (10mM Tris-HCL (pH 8.0), 100mM NaCl, 1mM EDTA, 0.5mM EGTA, 0.1% sodium Deoxycholate) (58) followed by 4°C sonication at an amplitude of 100 for 10 cycles of 30 second pulses and 30 second pauses. Sonicated samples were then centrifuged at 14,000rpm for 10 minutes at 4°C. A small aliquot of supernatant was saved for input in ChIP sequencing. 100µl/sample of Protein A Dynabeads (Thermo Scientific # 10001D) were incubated with either 0.66µg of HA (Cell Signaling Technology #3724S) or normal rabbit IgG (Millipore #12-370) antibody for 10 minutes at 4°C with rotation including the appropriate wash steps as described in the manufacturer’s protocol. The cell lysate’s supernatant protein concentration was measured using the Pierce BCA protein assay. A volume equivalent to 500µg of protein in each supernatant was incubated with the Dynabeads overnight at 4°C with rotation. Following incubation, Dynabeads were washed six times with RIPA buffer (150mM NaCl, 10mM Tris (pH 7.2), 0.1% SDS, 1% Triton X-100, 1% sodium Deoxycholate) utilizing a magnetic stand for separation of beads and buffer. The beads were then washed once with TE buffer (10mM EDTA, 50mM Tris-HCL (pH 7.4)). ChIP samples and inputs were de-crosslinked with 200µl of elution buffer (1% SDS, 0.1M NaHCO3) overnight at 65°C. DNA was purified using the QIAquick PCR purification kit following the manufacturer’s protocol. ChIP qRT-PCR quality control for *GREB1* was performed. HA ChIP was performed three times independently. The first biological replicate of samples was sent to the Health Sciences Sequencing Core at Children’s Hospital of Pittsburgh (HSSC, Pittsburgh, Pennsylvania) for TruSeq ChIP Library preparation (Illumina) using paired indexing. Qubit quantification and TapeStation HS D1000 was performed for library quantity control. Additional size fragment purification was performed by HSSC with the remaining biological replicates. DNA sequencing was performed on an Illumina NextSeq 2000 P3 flow cell 100 at 2×61bp reads with approximately 80 million reads per sample (overall 1.2 billion reads). ChIP sequencing data is in the process of being deposited in the Gene Expression Omnibus database.

ChIP-seq reads were aligned to Hg38 reference genome assembly using Bowtie 2.0 (59), and peaks were called using MACS2.0 with a q-value<0.1 (60) and visualized with the WashU Browser (61). We used the Diffbind (62) package identify differentially expressed binding sites and analyze intersection ratios with other datasets. Briefly, all BED files for each cell line were merged and binding intensity was estimated at each site based on the normalized read counts in the BAM files. Pairwise comparisons between ESR1-WT and fusion samples were performed to calculate FC. Heatmaps and intensity plots for binding peaks were visualized by Deeptools (63). For gene annotation from ER binding sites, gained ER peaks were selected and annotated genes were inspected in a ± 2kb range of the ER peaks using ChIPseeker (64).

### Cell growth assay

Hormone deprived T-47D shESR1 and MDA-MB-134-IV shESR1 cells were seeded in replicates of six in 2D (Fisher #353072) and 3D (ultra-low attachment (ULA), Corning #3474) 96-well-plates at a cell density of 6,000 cells/well and 8,000 cells/well, respectively. A day zero plate was seeded for 2D and 3D conditions with corresponding day 4, approximate day 7 and approximate day 14 plates for each treatment condition. 24-hour post-seeding cells were treated with either 1nM estradiol, 100nM ICI, a combination of both, or equivalent volume of 100% ethanol. Plates were collected on day 0, day 4, at one week and at two weeks and measured by CellTiter-Glo (Promega #PR-G7573) following the manufacturer’s protocol. Cell viability values were calculated by blank cell deductions and normalization to corresponding day zero readings.

### Colony formation assay

Hormone deprived T-47D shESR1 and MDA-MB-134-IV shESR1 cells were seeded in triplicates in 6-well plates (Fisher #08-772-1B) at a cell density of 36,000 cells/well and 70,000 cells/well, respectively. Cells were monitored daily and IMEM medium enriched with 10% CSS was refreshed every four to five days. Assay was stopped when a well displayed multiple overlapping individual colonies. Cells were fixed on ice with ice cold 100% methanol and stained with 0.5% Crystal Violet (Sigma-Aldrich #C0775) in 40% methanol. Wells were imaged on an Olympus SZX16 dissecting microscope. Quantification was performed utilizing the ImageJ threshold counting tool and particle analyzer plugin.

### Wound scratch assay

Hormone deprived T-47D shESR1 and MDA-MB-134-IV shESR1 cells were seeded in replicates of 8 onto Imagelock 96-well plates (Essen Bioscience #4379) pre-coated with Matrigel (Corning #356237) at cell densities of 150,000 cells/well and 250,000 cells/well, respectively. 24 hours post seeding wounds were made to the middle of each well using a Wound Maker (Essen Bioscience #4493). T-47D shESR1 cells were washed twice with 100µl 1X PBS, MDA-MB-134-IV shESR1 cells were washed once with 100µl 1X PBS. Desired treatment, supplemented with 5µg/ml Mitomycin C (Sigma Aldrich #10107409001), was added to cells. Cells were monitored and wound closure was imaged every four hours over the course of 72 hours using the IncuCyte Imaging system (Sartorius, Albuquerque, New Mexico). Wound closure density was calculated using the IncuCyte system’s wound scratch analysis module.

### Statistical Analysis

GraphPad Prism software version 7 and R version 4.0.0 were used for statistical analysis. All experimental results included biological replicates and were shown as a representative reading ± SEM. 2way ANOVA with posthoc Dunnett’s multiple comparisons test showing enrichment in comparison to corresponding ESR1-WT treatment (#) or to intra-construct vehicle treatment (*) in each assay. */#p=0.0332, **/##p=0.0021, ***/###p=0.0002, ****/####p<0.0001.

## Authors’ Contributions

**M.E. Yates:** conceptualization, funding acquisition, data curation, formal analysis, investigation, visualization, methodology, project administration, writing-original draft, writing-review and editing. **Z. Li**: data curation, formal analysis, investigation, visualization, methodology, writing-review and editing. **Y. Li:** data curation, formal analysis, writing-review and editing. **H. Guzolik:** data curation, formal analysis, investigation, visualization, methodology, writing-review and editing. **X.S. Wang:** investigation, resources, writing-review and editing. **T. Liu:** conceptualization, data curation, formal analysis, investigation, visualization, methodology, writing-review and editing. **J. Hooda:** conceptualization, data curation, investigation, resources, supervision, project administration, writing-review and editing. **J.M. Atkinson:** conceptualization, resources, supervision, project administration, writing-review and editing. **A.V. Lee:** conceptualization, funding acquisition, resources, methodology, supervision, project administration, writing-review and editing. **S. Oesterreich:** conceptualization, funding acquisition, resources, methodology, supervision, project administration, writing-review and editing.

## Acknowledgments

The authors would like to thank the numerous foundations that have supported our work. In addition, we would like to acknowledge Dr. Matthew Ellis for his generous gift of the ESR1-DAB2 plasmid. This research was supported in part by the University of Pittsburgh Center for Research Computing, RRID:SCR_022735, through the resources provided. Specifically, this work used the HTC cluster, which is supported by NIH award number S10OD028483. This project is funded, in part, under a Grant with the Pennsylvania Department of Health. The Department specifically disclaims responsibility for any analyses, interpretations, or conclusions. In addition, this work was supported by the National Cancer Institute (F30CA250167 to M.E.Y., R01CA256161 to A.V.L. & S.O., R01CA181368 to X-S.W., R01CA183976 to X-S.W., R21CA237964 to X-S.W.), the HCC Cancer Biology Program Pilot Funding (X-S.W.), the Shear Family Foundation (A.V.L., S.O. & X-S.W.), the CCSG Dev Funds (P30CA047904-31 to X-S.W.), the Hillman Foundation (X-S.W.), the PA Breast Cancer Coalition (A.V.L.) and the Breast Cancer Alliance (S.O.). Lastly, we would like to acknowledge that Figure 7 and Supplementary Figure S1 were created with BioRender.com.

## Supplementary Figure Legends

**Supplementary Figure S1:**
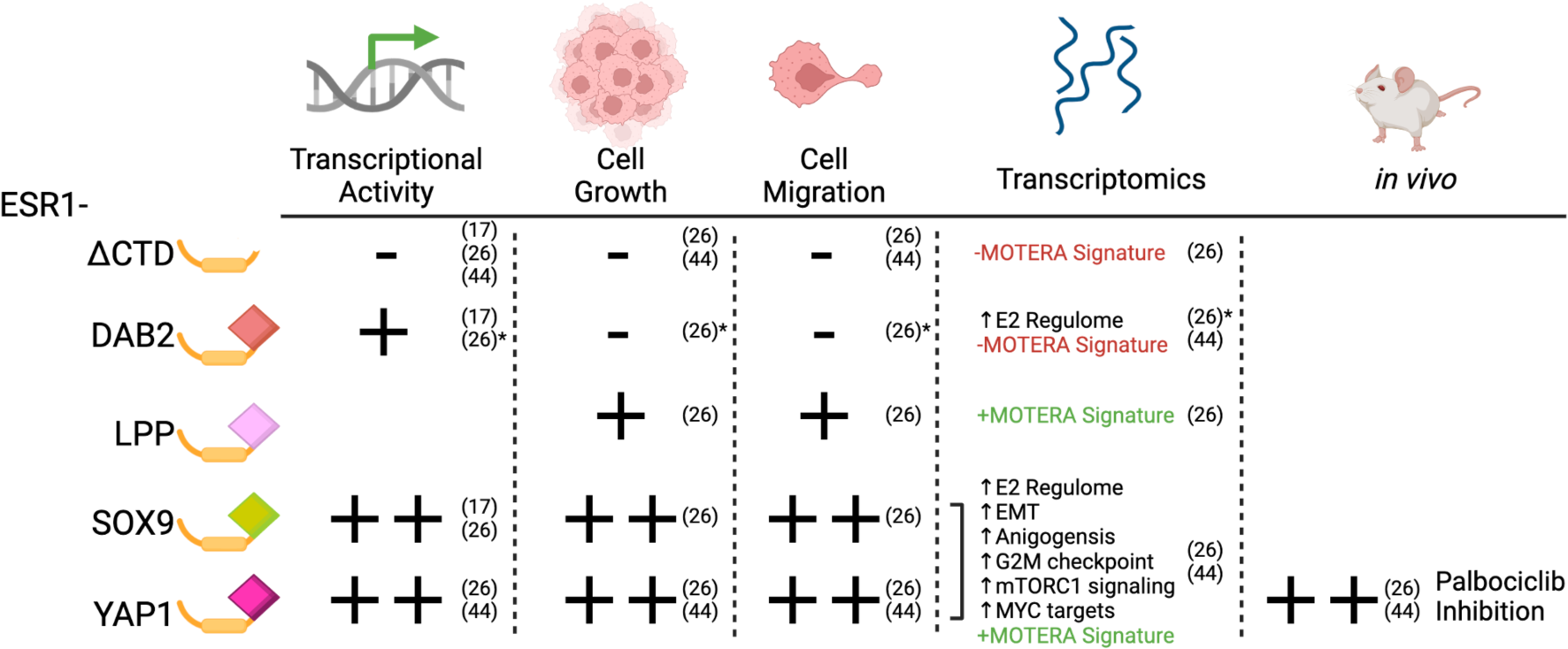
Schematic of previous ER fusion functional activity. Asterisks denote cell line dependent phenotypes. References reported in paratheses. Plus sign indicates reported activity of fusion in corresponding manuscript. Minus sign signifies no appreciable activity.

**Supplementary Figure S2:**
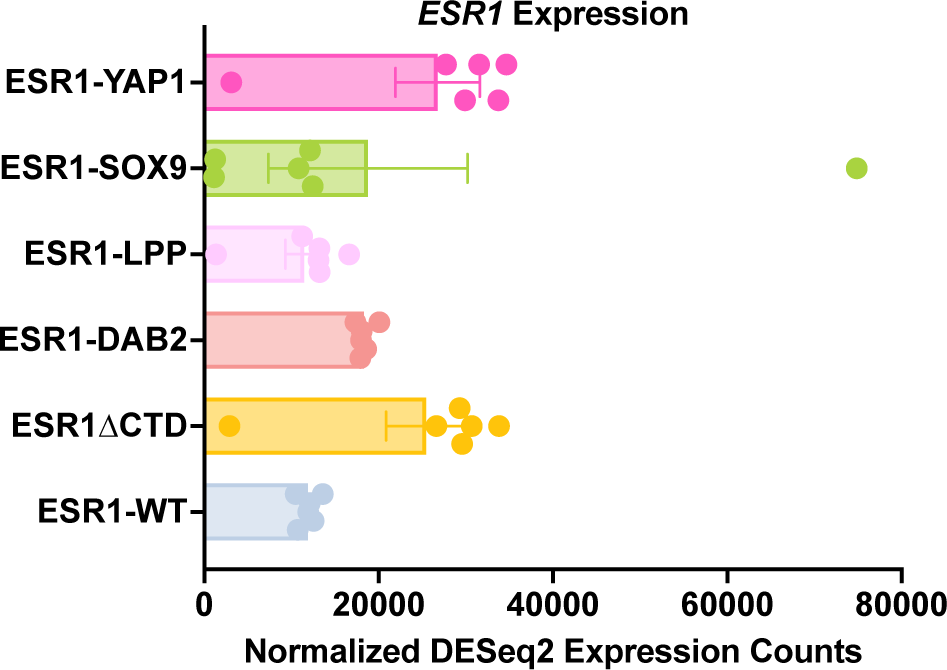
*ESR1* RNA-seq expression across cell lines. *ESR1* expression from normalized DESeq2 values.

**Supplementary Figure S3:**
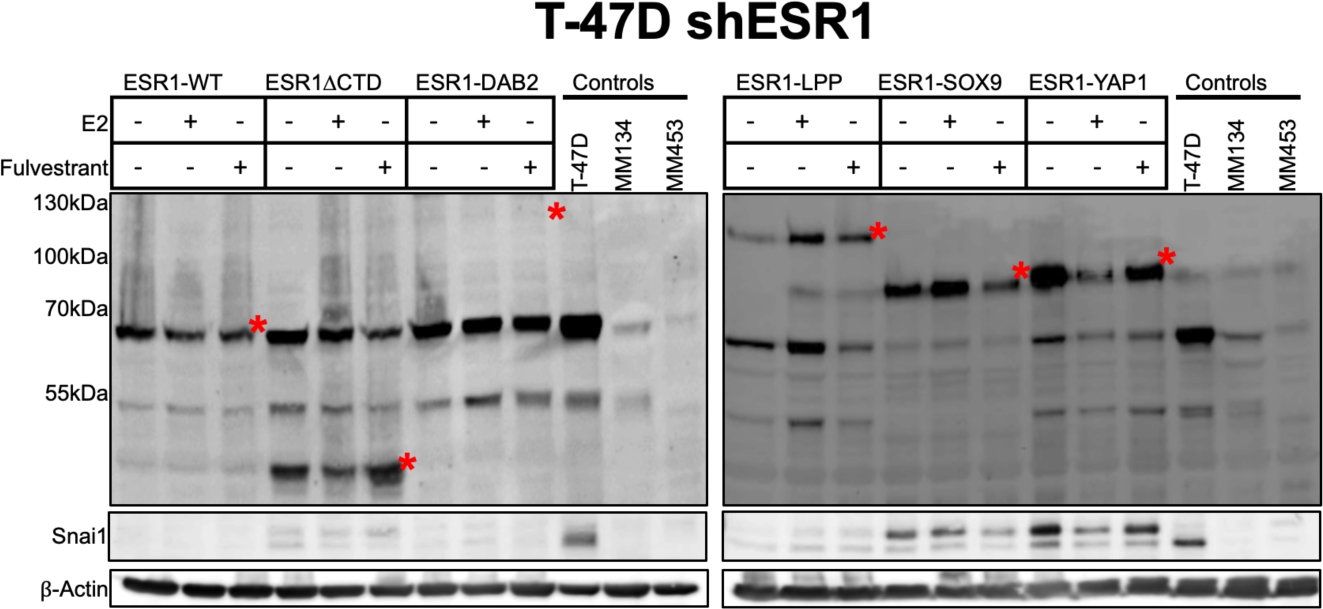
NST fusion positive cells express SNAI1. Anti-ER and Anti-Snai1 immunoblots of stably overexpressed constructs in T-47D cell lines. Hormone deprived cells were stimulated with either control, 1nM E2 or 100nM ICI. Construct proteins denoted by red asterisks in all blots. β-actin serves as loading control.

**Supplementary Figure S4:**
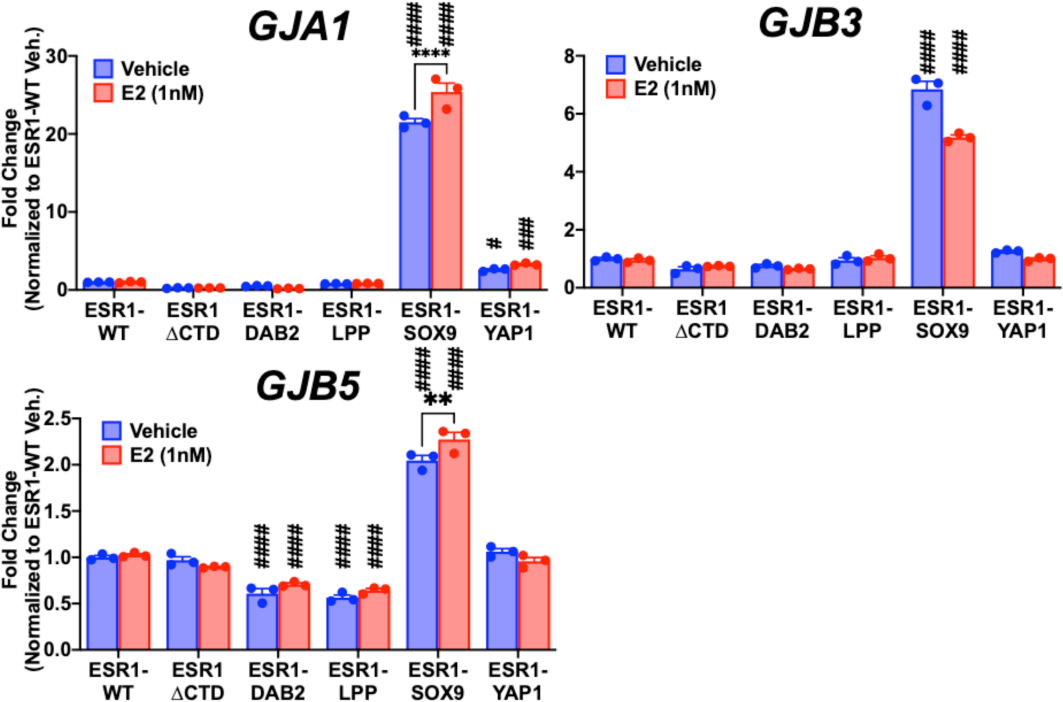
ESR1-SOX9 enriched in gap junction gene family. qRT-PCR in T-47D shESR1 cellular RNA targeting gap junction genes *GJA1*, *GJB3*, and *GJB5*. 2way ANOVA with posthoc Dunnett’s multiple comparisons test showing enrichment in comparison to ESR1-WT (#) or to intra-construct vehicle treatment (*). Representative experiment shown with bar graph equivalent to readings ± SEM, n=3 for each experiment. */#p=0.0332, **/##p=0.0021, ***/###p=0.0002, ****/####p<0.0001.

**Supplementary Figure S5:**
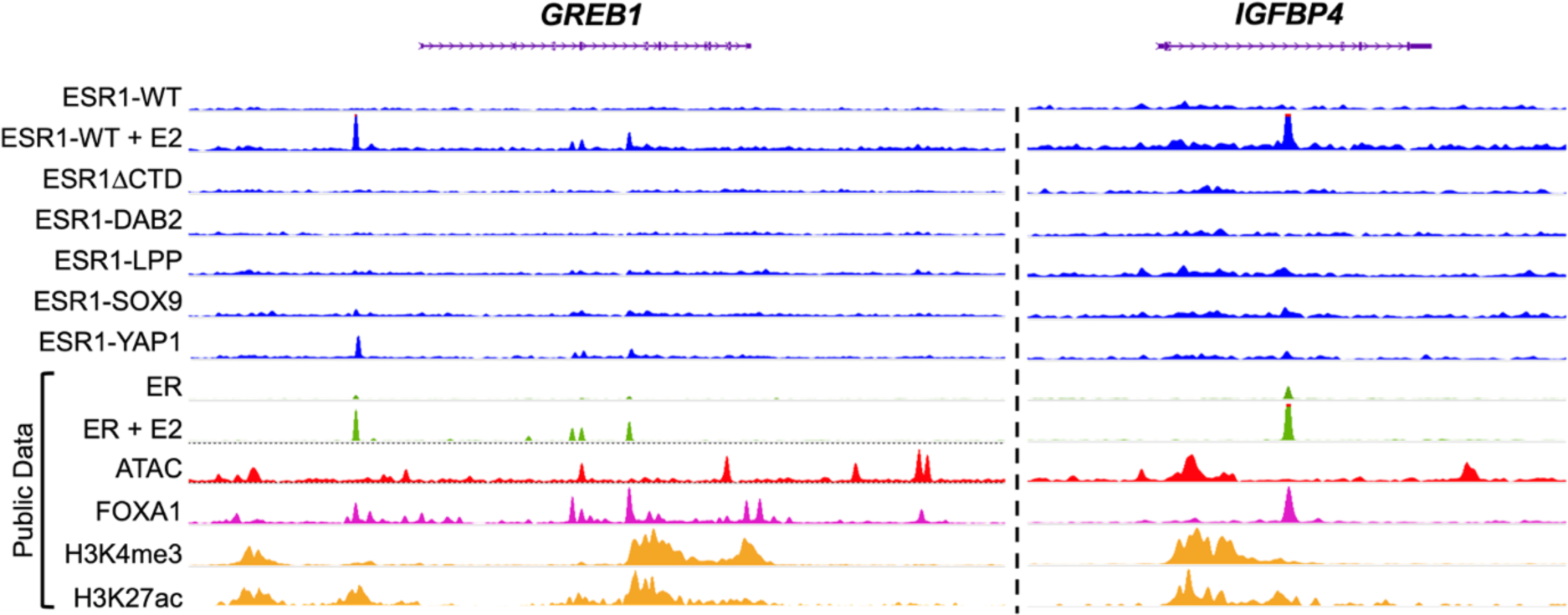
HA ChIP-seq fusion construct binding peaks at *GREB1* and *IGFBP4* loci. Binding enrichment of HA tagged wildtype ER stimulated with or without E2, HA tagged truncated ER, and HA tagged fusion constructs at *GREB1* and *IGFBP4* loci utilizing HA ChIP-seq data. Relative peak enrichment compared to publicly available ChIP-seq binding of ER, ER stimulated with E2, ATAC, FOXA1, H3K4me3, and H3K27ac at *GREB1* and *IGFBP4*.

## Supplementary Table Legend

**Supplementary Table S1:** ERE enrichment in HA ChIP-seq derived T-47D shESR1 fusion cistromes. Find Individual Motif Occurrences (FIMO) analysis for *ESR1* motif enrichment in HA ChIP-seq data of control and fusion constructs in T-47D shESR1 cell lines.

|  | ESR1 motif | Gained Peaks | Ratio<br>(ER motif/gained peak) | Peak % containing ERE |
| --- | --- | --- | --- | --- |
| ESR1-WT + E2 | 986 | 748 | 1.318181818 | 76.87% |
| ESR1ΔCTD | 2 | 88 | 0.022727273 | 2.270% |
| ESR1-DAB2 | 1 | 105 | 0.00952381 | 0.100% |
| ESR1-LPP | 57 | 117 | 0.487179487 | 18.80% |
| ESR1-SOX9 | 31 | 17 | 1.823529412 | 94.12% |
| ESR1-YAP1 | 546 | 1040 | 0.525 | 23.85% |

